# Modelling resource-driven movements of livestock herds to predict the impact of climate change on network dynamics

**DOI:** 10.1101/2024.05.09.593362

**Authors:** Tijani A. Sulaimon, Divine Ekwem, Douglas Finch, Paul I. Palmer, Sarah Cleaveland, Jessica Enright, Paul C. D. Johnson, Rowland Kao

**Affiliations:** The Roslin Institute, University of Edinburgh, Easter Bush Campus, Midlothian, United Kingdom; Royal (Dick) School of Veterinary Studies, University of Edinburgh, Easter Bush Campus, Midlothian, United Kingdom; School of Biodiversity, One Health and Veterinary Medicine, University of Glasgow, Glasgow, United Kingdom; International Livestock Research Institute, Nairobi, Kenya; School of GeoSciences, University of Edinburgh, Edinburgh, United Kingdom; School of Computing Science, University of Glasgow, Glasgow, United Kingdom; School of Physics and Astronomy, University of Edinburgh, Edinburgh, United Kingdom

**Keywords:** Network model, Livestock movement, Agropastoral, Climate change, Tanzania

## Abstract

In East Africa, climate change is likely to profoundly impact livestock management and the potential spread of infectious diseases. Here, we developed a network model to describe livestock movements to grazing and watering sites, fitted it to data from the Serengeti district of Tanzania, and used it to explore how projected changes in resource availability due to climate change could impact future network structures and therefore infectious disease risks, using 2050 and 2080 as exemplar scenarios. Our modelled networks show increased connections between villages in grazing and watering networks, with connectivity increasing further in the future in correspondence with changes in vegetation and water availability. Our analyses show that targeted interventions to efficiently control regional disease spread may become more difficult, as village connectivity increases and disease vulnerability becomes more evenly distributed. This analysis also provides proof of principle for a novel approach applicable to agropastoral settings across many developing countries, where livestock trade plays a crucial role in maintaining local livelihoods but also in spreading disease.

## 1 Introduction

Climate change is likely to have a profound impact on the way livestock in East Africa are managed. Livestock play a critical role in the livelihoods of many households in rural Africa [1], providing security for households practising other agricultural systems with unpredictable yields, such as crop production [2]. This security can be compromised by infectious diseases that threaten livestock production, health, and overall welfare [3]. Thus, understanding how management changes could impact infectious disease transmission is critically important, but until now, it remains largely understudied. A key consideration is livestock movements, which are known to drive the transmission and spread of infectious diseases within and between populations [4]. These movements are mainly driven by trade activities or the need to access resources to ensure survival [5].

As the demand for livestock and animal products increases due to population growth and urbanisation [6], livestock production is facing significant and ongoing challenges from a variety of factors, including climate, land use, and policy changes [7–9]. Environmental and climatic changes can alter resource abundance and alter land fragmentation thereby influencing livestock contact patterns and with uncertain impact on disease transmission [6, 10].

Quantifying this impact requires models for livestock movement patterns that consider the impact of resources constraints. Most methods for quantifying livestock movement patterns rely on intensive data collection [11, 12], augmented by statistical modelling approaches [13–15]. Recently, network simulation modelling approaches have greatly facilitated the generation and evaluation of contact patterns between animal populations.

These network simulation approaches have been successful at describing livestock trade movements [16–18]. However, little attention has been paid to resource-driven movements of livestock herds in agropastoral settings [19]. This study aims to develop a spatially explicit probabilistic model for herd movements to shared resources that can be used to suggest future changes in network structure due to the impact of changing resource availability.

Here, we show that a simulated movement model of contacts between villages generated based on distance and abundance of resources (Section 4.2) shows good fidelity to recorded patterns of cattle movements. We used the Normalised Difference Vegetation Index (NDVI), which is a measure of vegetation greenness [20], as an indicator of resource abundance and availability in different seasons. It also represents the attractiveness of various resource locations [21], as wet and green pasturelands, which are more likely to be proximal to a water source, are usually preferred to dry areas [22, 23]. We focused on contacts at communal livestock resource areas (grazing and watering points) in an agropastoralist setting, where movements occur daily. Based on the summary statistics of the network of connections between 95 villages through the use of communal resources in the Serengeti district ([15]), we used approximate Bayesian computation (ABC) to estimate the parameters of the movement model (Section 4.4). We evaluated the performance of the model by comparing the networks generated from the posterior predictive distribution of the fitted movement model with the observed networks (Sections 2.1 and 2.2). Finally, we used the fitted model to explore the long-term impacts of a range of future scenarios resulting from climate and policy change through removal of or change in resource availability (Section 2.4).

## 2 Results

We developed a kernel-based movement model, outlined in detail in section 4, to simulate monthly networks of village-to-village contacts facilitated by communal grazing and watering points. We fitted our model to the observed network data using ABC. Here, we report on the posterior distributions of fitted model parameters, model performance, and projected changes in network structures derived from our model predictions.

### 2.1 Posterior distributions

The posterior medians and 95% credible intervals for all parameter estimates are presented in Figures A4 and A5, along with their corresponding priors. The parameters of the grazing and watering networks showed considerable heterogeneity in the movement distances and the mean number of movements over months, highlighting the importance of modelling them separately. The distance effect varies between seasons, with a higher median value during the wet season compared to the dry seasons in both the watering and the grazing networks. In general, the posterior distributions of the distance and the mean parameter in the negative binomial distribution are narrow, while those of the NDVI effect and the dispersion parameter show wide intervals. Estimates of the NDVI effect show considerable uncertainty, likely due to the similarity of NDVI values at grazing and watering sites, leading to wider distributions across all months and seasons in both networks. The mean parameter of the negative binomial distribution is lower in dynamic month networks compared to static season networks. Note that inferences on the distance effect are sensitive to prior values (see prior distributions in Table A3), and the prior values for the mean parameter were restricted based on the number of links in each network. Setting a broader prior range for monthly networks led to an unrealistic number of links at the annual timescale.

### 2.2 Model performance

To evaluate the predictive ability of our kernel model against observed data using network properties, we simulated 100 monthly bipartite networks, connecting villages to resource areas, for comparison with the properties of the observed networks. We transformed the bipartite networks into monthly village-village connections (see subsection 4.3). Generally, the degree distributions and number of unique links in the simulated monthly inter-village connections at shared grazing and watering points were consistent with the observed monthly networks. Figure 1 shows how the monthly simulated networks closely match the observed networks in terms of the number of unique links in both the grazing and the watering networks, with a slight overestimation in January and a slight underestimation in the drought months of the grazing networks (Figure 1).

**Fig. 1:**
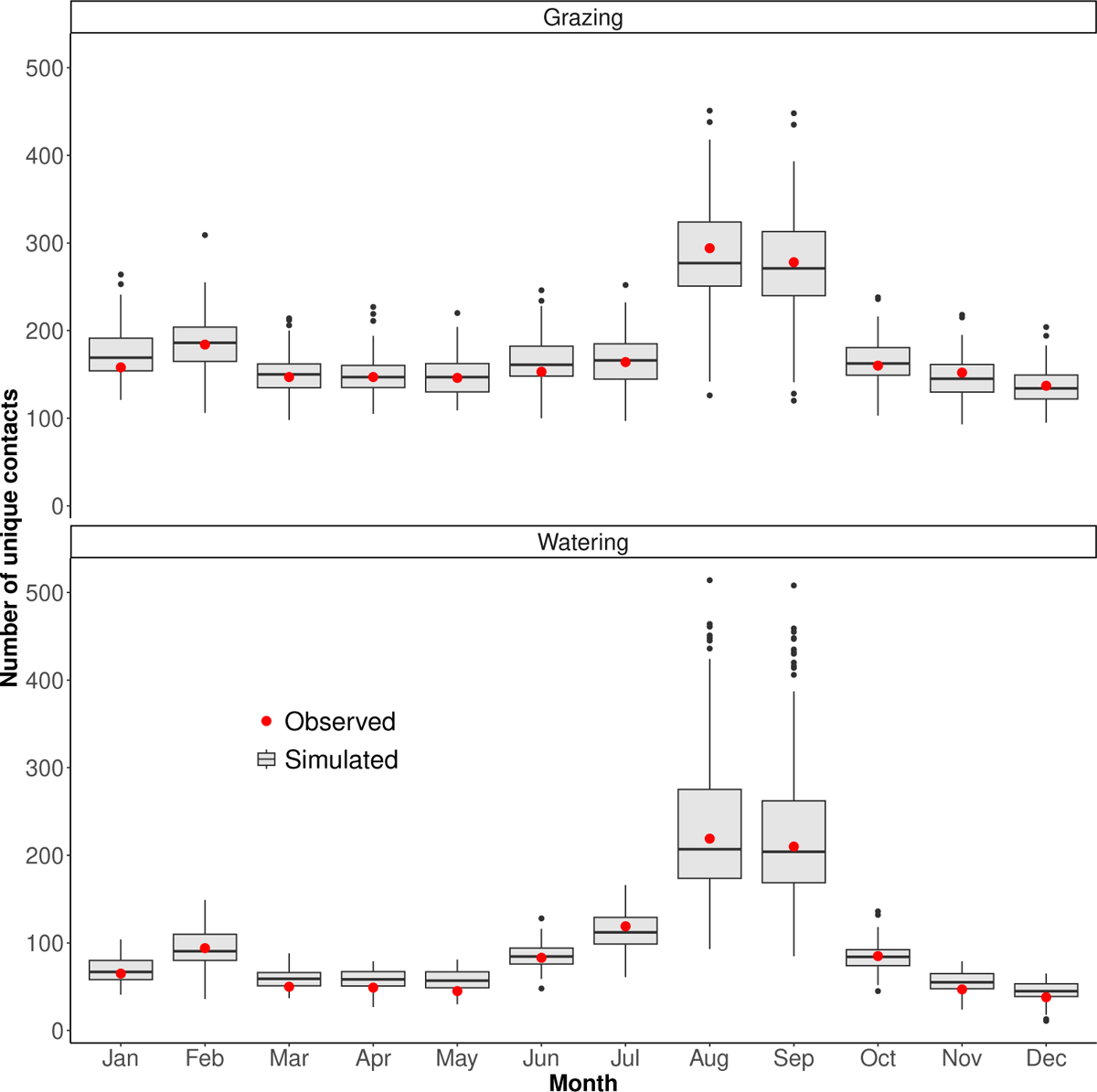
Comparison of the distribution of the number of unique contacts between 100 networks simulated from the movement model (box plots) and the observed network (red points).

The degree distribution of the monthly simulated grazing and watering networks was in good agreement with the corresponding degree distributions of the observed monthly grazing and watering networks (Figure 2). The observed degree distribution fell within the simulated degree distributions for most ranges, but with minor deviations in the dry months. At an annual scale, the aggregated monthly simulated networks were notably similar to the observed annual network, with most of the measured metrics in the observed network lying well within the range of simulated values (Table 1). However, there was some loss of fidelity found in the pairwise geographic distance between connected villages across all months in both networks, which were consistently underestimated (Figure 3).

**Fig. 2:**
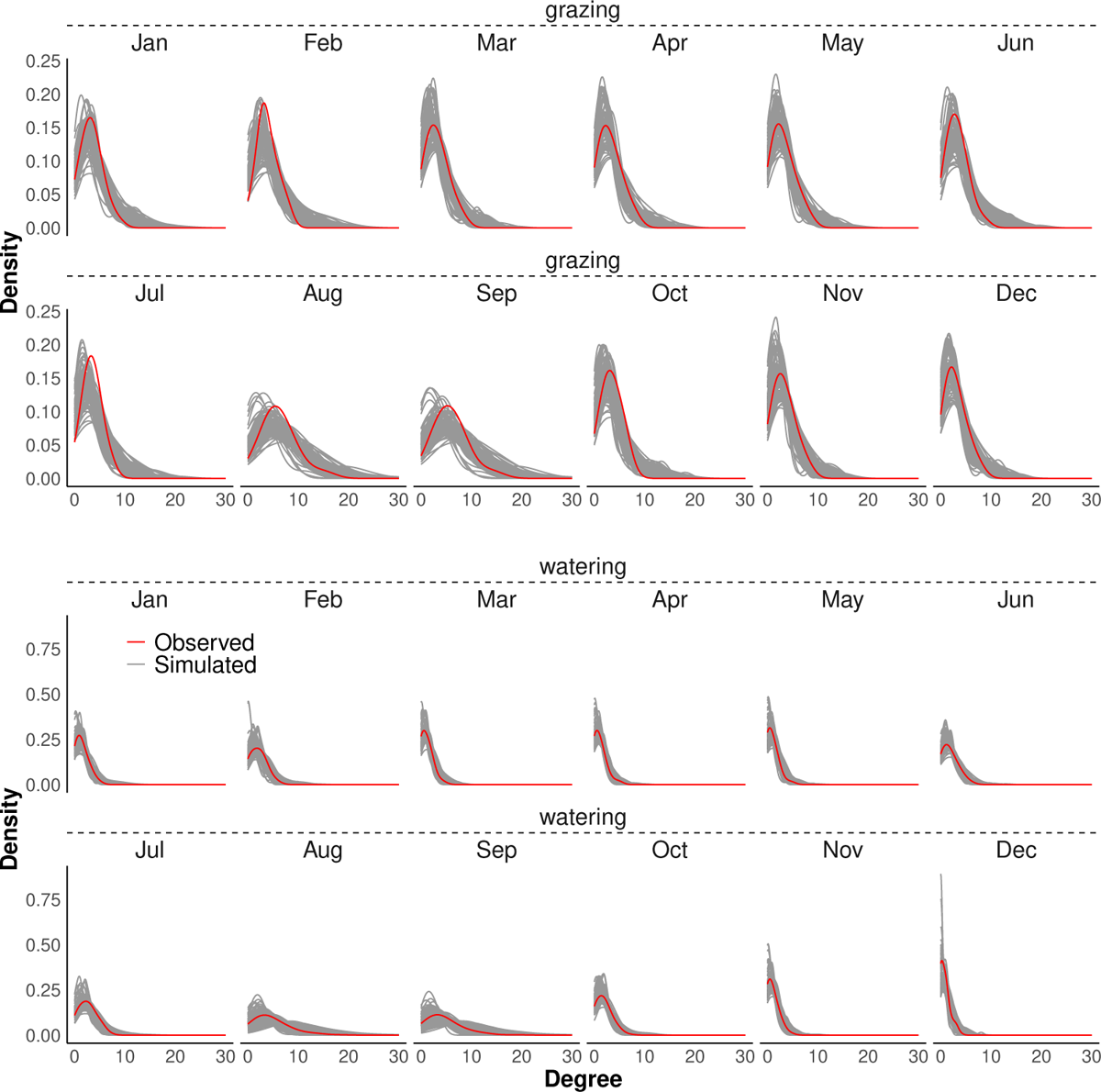
Comparing the degree centrality scores of monthly grazing and watering networks. The grey lines represent degree distributions of monthly networks from 100 simulations of the kernel movement model. In contrast, the red lines show the observed monthly contact networks from observed data.

**Fig. 3:**
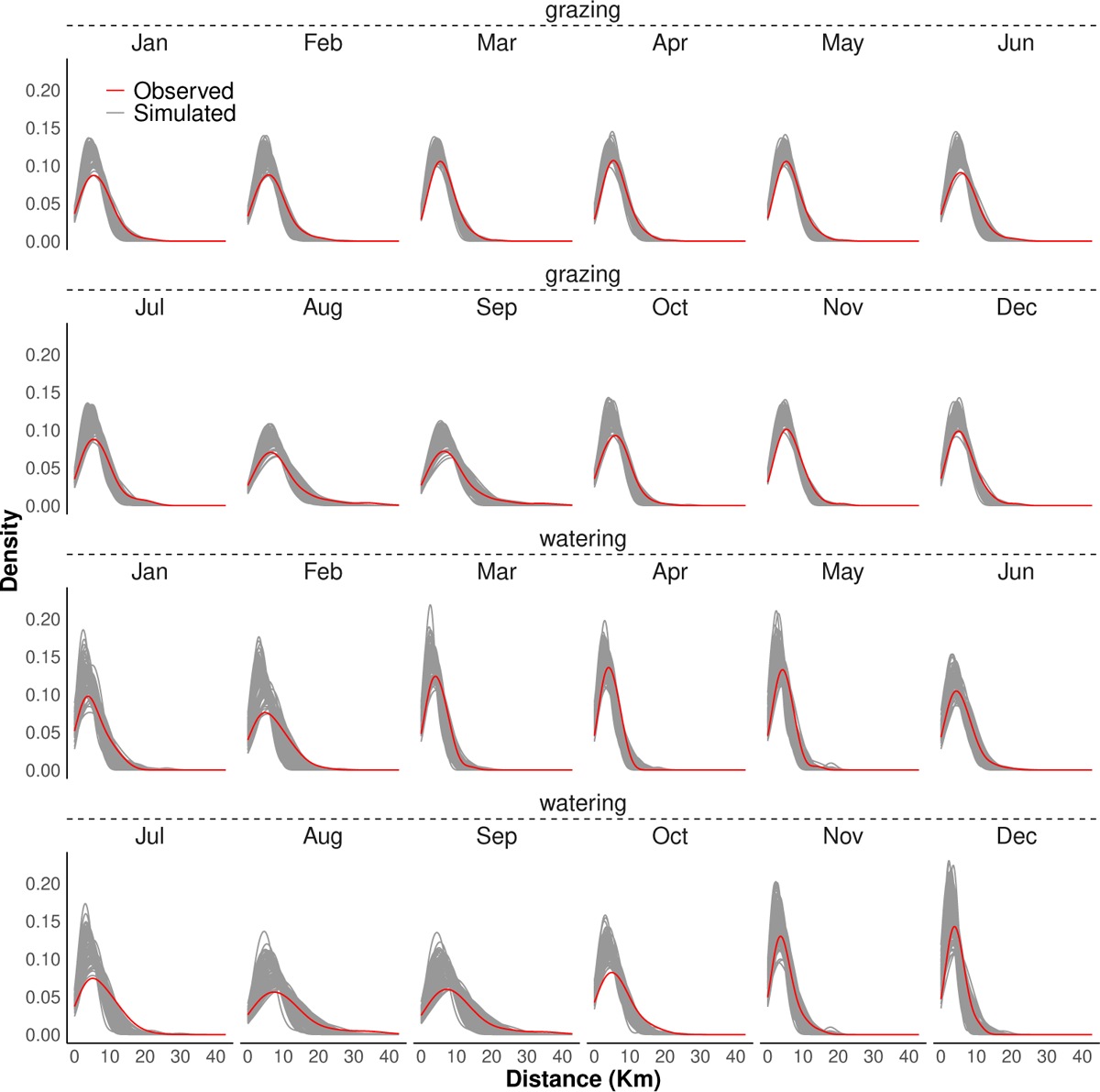
Distribution of monthly distance between connected villages from 100 simulated (gray) and observed (red) networks.

**Table 1:** Observed global and summary node-level statistics of year networks compared to the mean and standard deviations (sd) of 100 simulated grazing and watering networks.

| Measure | Grazing networks |  |  | Watering networks |  |  |
| --- | --- | --- | --- | --- | --- | --- |
|  | Observed Value | Simulated Mean | sd | Observed Value | Simulated Mean | sd |
| Number of active nodes | 95 | 94.9 | 0.24 | 91 | 94.8 | 0.51 |
| Number of distinct links | 353 | 489 | 41.9 | 246 | 384 | 85.1 |
| Mean degree | 7.43 | 10.3 | 0.88 | 5.41 | 8.09 | 1.79 |
| Maximum degree | 19 | 24.8 | 2.62 | 19 | 20.1 | 4.34 |
| Mean betweenness | 112 | 117 | 14.5 | 104 | 138 | 42.7 |
| Maximum betweenness | 983 | 1170 | 338 | 1100 | 1190 | 473 |
| Mean eigenvector | 0.07 | 0.06 | 0.002 | 0.06 | 0.06 | 0.005 |
| Maximum eigenvector | 0.30 | 0.27 | 0.011 | 0.34 | 0.29 | 0.023 |
| Network diameter | 8.00 | 8.11 | 1.18 | 10 | 9.07 | 2.64 |
| Mean network distance | 3.38 | 3.48 | 0.31 | 3.78 | 3.99 | 0.90 |
| Transitivity | 0.48 | 0.57 | 0.020 | 0.51 | 0.47 | 0.033 |
| Assortativity | 0.34 | 0.51 | 0.059 | 0.39 | 0.38 | 0.12 |

### 2.3 Predicted future NDVI

The historical and projected mean temperature, precipitation, and NDVI trends from 1982 to 2100 are shown in Figure B11. According to the SSP585 scenario (Shared Socioeconomic Pathway 5, which represents the high end of the range of future emission pathways), the Serengeti district is expected to experience sustained increases in both mean temperature and precipitation over this entire period [24]. Additionally, Figure B12 shows the seasonality in mean temperature, precipitation, and predicted NDVI for each year from 2015 to 2100, with consistent patterns observed throughout these years. Precipitation peaks during the long wet season (March to May) and reaches its lowest during the long dry season (July to September). This pattern mirrored the predicted NDVI, which also showed its lowest values in August and September.

In Figure 5, it is observed that during the wet season (April), all areas have NDVI values greater than 0.45 in 2016, 2050, and 2080, indicating the availability of resources during these periods. However, during the long dry season (August), the northern part has higher NDVI values compared to the southern part in 2016, with greener areas predominantly concentrated in the northwest. By 2050, many areas will experience reduced predicted NDVI values, with a more pronounced reduction by 2080.

**Fig. 4:**
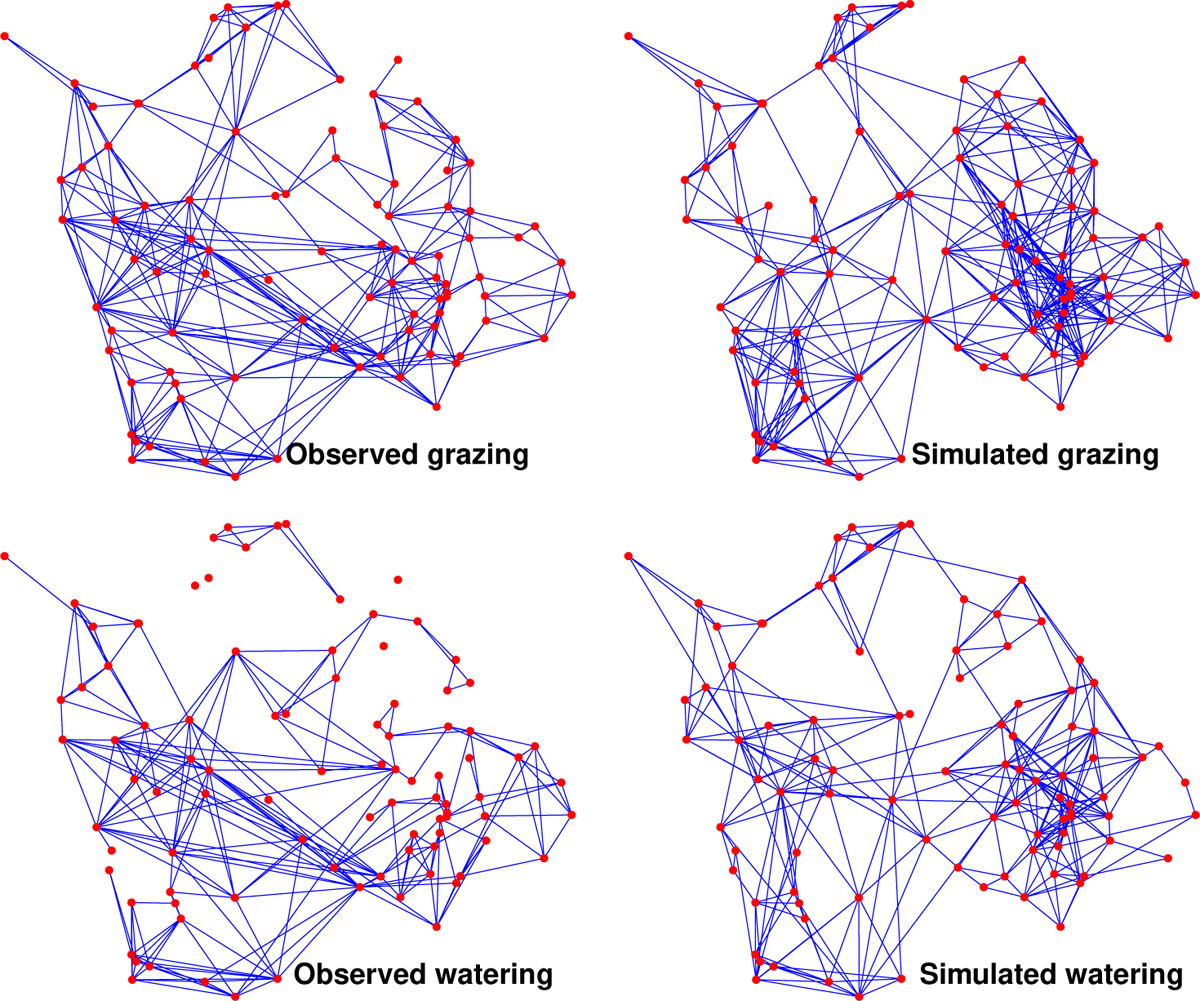
Visual comparison of observed and simulated year networks. The simulated network is a single realisation sampled from 100 kernel-generated networks.

**Fig. 5:**
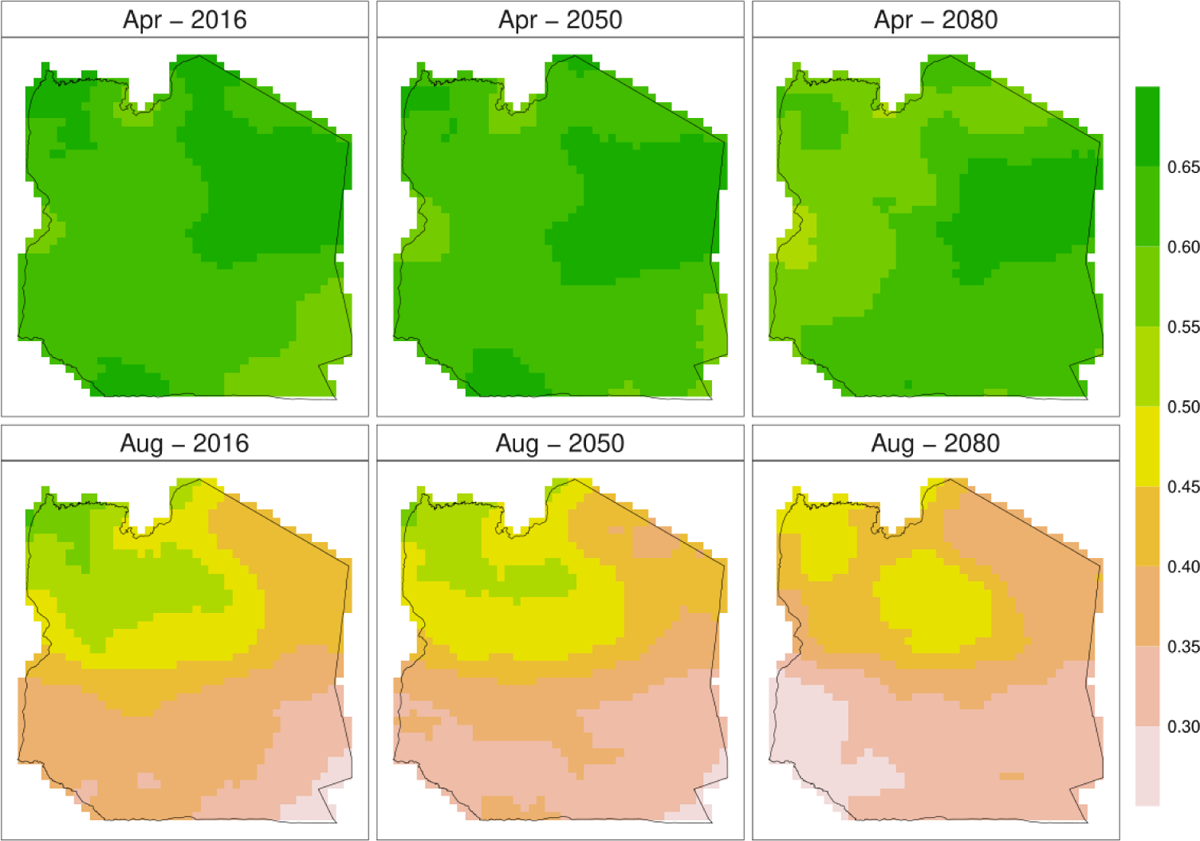
Current and projected NDVI distribution in wet and dry season months. Regions with NDVI *>* 0.45 are suitable for grazing and watering, while those below are unsuitable.

### 2.4 Future changes in network structures

To evaluate future changes in network structures for both grazing and watering networks, we examined exemplar scenarios for 2050 and 2080, using the simulated values of 2016 in Table 1 as a baseline for comparison. Additionally, we investigated the effects of a 10% reduction in grazing and watering points for both 2050 and 2080. The changes in summary statistics from 100 simulated future networks are shown in Figure 6.

**Fig. 6:**
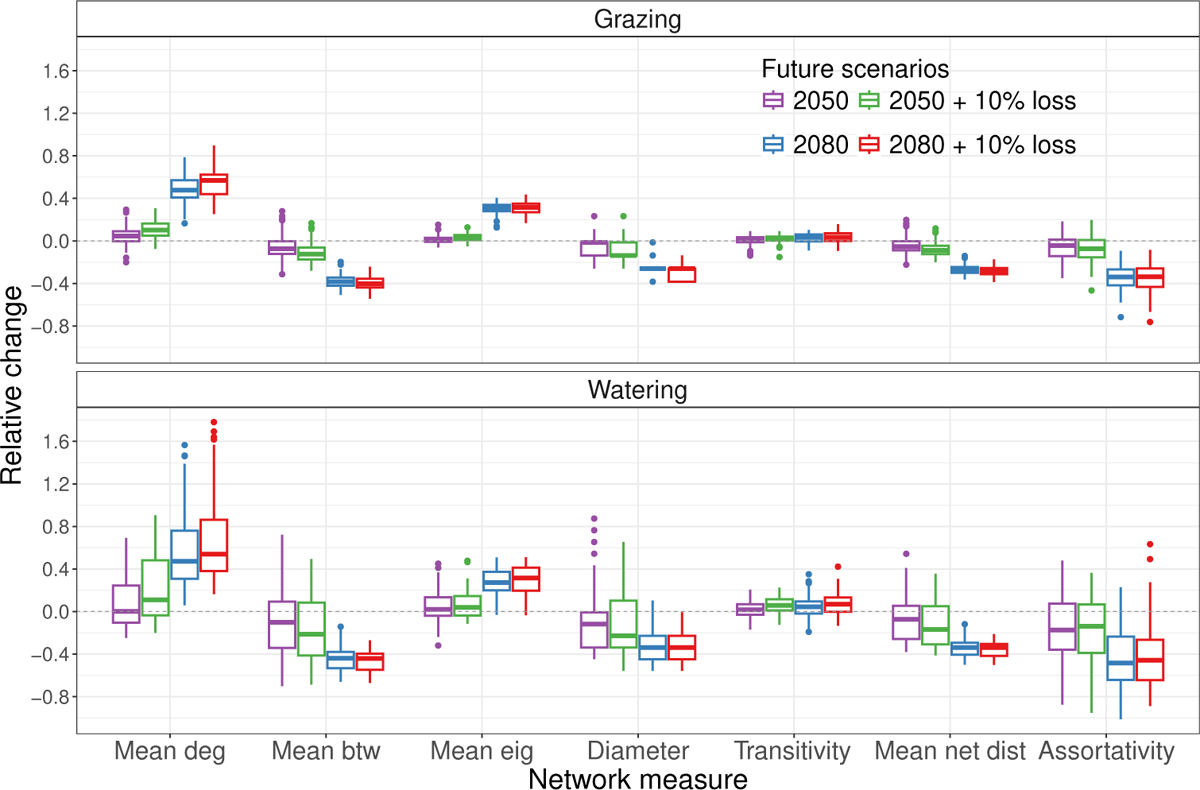
Changes in global and summary node-level statistics of simulated future projected year networks (2050 and 2080) relative to the summary statistics of the baseline simulated networks (2016). The 2016 summary statistics represent simulated values evaluated against observed summary statistics in Table 1. The scenarios for 2050 and 2080 are future projections, with and without a 10% loss in grazing and watering sites. Deg degree; Btw betweenness; Eig eigenvector; Net dist network distance.

The grazing and watering networks have a similar number of active nodes (villages), with each village maintaining at least one annual contact with other villages in all scenarios. The number of links is projected to increase in future scenarios for both networks, with a more pronounced increase observed in the 2080 scenarios. In the grazing network, the number of links increased by 4.6% (IQR = [0%, 9.0%]) in 2050 and by 47.8% (IQR = [40.9%, 57.1%]) in 2080 compared to the 2016 scenario. Similarly, in the watering network, the number of links increased by 0.2% (IQR = [*−*10.5%, 24.4%]) and 47.3% (IQR = [30.8%, 76.0%]) in 2050 and 2080, respectively. With a 10% reduction in grazing and watering points, the number of links in the grazing network increased by 10.2% (IQR = [5.03%, 16.4%]) and 56.9% (IQR = [44.1%, 62.4%]), and by 11.0% (IQR = [*−*3.66%, 48.2%]) and 54.0% (IQR = [38.0%, 86.3%]) in the watering networks in 2050 and 2080. The mean degree followed a trend similar to the number of links in both networks.

Mean betweenness values showed a decreasing trend in both networks. In 2050, mean betweenness decreased by 7.32% (IQR = [*−*12.2%*, −*0.3%]) in the grazing network and by 10.2% (IQR = [*−*34.2%, 9.27%]) in the watering network, further decreasing to 38.4% (IQR = [*−*42.1%*, −*34.5%]) and 44.0% (IQR = [*−*53.3%*, −*38.0%]) in 2080. In scenarios where there was a 10% reduction in resource points, the mean betweenness in the grazing and watering networks decreased by 12.5% (IQR = [*−*17.4%*, −*6.21%]) and 21.3% (IQR = [*−*41.3%, 8.32%]) in 2050 and by 40.2% (IQR = [*−*43.8%*, −*35.6%]) and 44.1% (IQR = [*−*54.8%*, −*39.7%]) in 2080, respectively.

In grazing and watering networks, the relative change in mean eigenvector centrality values was 0.83% (IQR = [*−*0.92%, 2.95%]) and 2.06% (IQR = [*−*3.95%, 13.5%]) in 2050, and 31.1% (IQR = [28.0%, 33.8%]) and 27.3% (IQR = [20.0%, 37.3%]) in 2080 under the assumption of no loss of resource points. These changes are similar to those observed when we introduced a loss of 10% in resource points.

In the 2050 and 2080 scenarios, the values of network diameter, mean network distance, and assortativity index consistently decreased. However, network clustering (transitivity) values showed a slight increase in future scenarios for both networks, with no noticeable difference between 10% loss and no loss scenarios.

## 3 Discussion

Local livestock movements, driven by the need to access resources for survival, play an essential role in connecting populations but also can be important for transmitting diseases between them. The potential impact of climate change on the availability of these key resources in the coming decades could alter the direction and extent of resource-driven livestock movements. Understanding the spatial pattern of contacts between populations and how they might change is plays a key role in the assessment and management of current and future disease risks. Here, we developed a kernel-based resource-driven model to simulate village contact networks through shared resources. This model estimates the probability of herd movements from villages to resource areas based on distance and vegetation abundance, allowing us to investigate how these movements might change in response to future climate change. The model was structured so that the kernel parameters varied for each month and season and for grazing and watering movements.

First, we used location and remotely sensed vegetation data to simulate livestock movement networks in agropastoral settings in developing countries and showed that the kernel model generated monthly networks were similar to the observed data. The centrality measures of villages derived from grazing and watering networks varied between seasons. Like the observed data, and our model predicted fewer contacts between villages during the wet season compared to the dry season. As well, herds travelled short geographical distances in the wet season compared to the long dry season. Long-distance movements occur during the dry seasons when grazing and water sources are limited. This seasonal variation in movement patterns highlights the dynamic nature of animal movements in response to changing environmental conditions.

To project future patterns we considered the impact of changes in the seasonal pattern of movements caused by changes in precipitation, due to its influence on the availability of pasturelands and water. Previous studies have shown that the frequency of use of resource areas is related to resource availability, which in turn is influenced by season [5, 15]. While two distinct seasons, wet and dry, are generally recognised, rainfall patterns are becoming increasingly inconsistent and unpredictable, resulting in long periods without rain even in the wet seasons and extended dry seasons and droughts [9, 25]. Ekwem et al. [15] showed that the change in rainfall patterns drove unplanned livestock movements, especially during drought. As temperature and precipitation are predicted to increase in the future, and East Africa is more influenced than most other parts of the world [7–9, 25], we hypothesise that the stresses on resources induced by climate change are likely to result in an overall more connected livestock population, with implications for infectious disease risks and therefore human and animal health. We incorporated predicted future NDVI within our model (based on SSP585 from CMIP6, see Appendix B) to generate future networks from estimated parameters. We also investigated how network structures are affected by the reduced availability of resource points through random removal. We used 2050 and 2080 as examples of future scenarios.

Our results show that while the number of active villages in both the grazing and watering networks remains relatively stable compared to the baseline year (each village maintaining at least one annual contact with other villages), the number of links is projected to increase in the future scenarios, with a more pronounced rise observed in the 2080 scenarios. Given our assumption that the grazing and watering behaviour of livestock herds remains the same and the relationship between NDVI and climate factors remains unchanged, changes in future network metrics suggest that, as villages are projected to have more contacts in the future, they are more likely to be infected during outbreaks and are likely to play an important role in propagating infections to other areas. The overall changes in node-level centrality measures suggest that the most critical villages currently identifiable by network centrality measures for disease control may be less valuable targets for intervention in the future as overall connectivity increases and thus their central nature diminishes.

The anticipated changes in the mean distance and diameter of future networks have implications for the speed of transmission of infectious diseases between villages. The mean network distance refers to the mean pairwise shortest network distance between villages, while the diameter represents the longest shortest path between any two villages within the network [26]. With projected reductions in both metrics in the future, diseases can spread between villages more rapidly.

Improvements to our kernel model would be necessary to reproduce the pattern of long-distance contacts present in the observed networks. This is likely due to more complex characteristics specific to resource locations, particularly in cases where proximal villages did not have the expected level of contacts, possibly due to communal or historical disputes/conflicts. Additionally, the absence of information on grazing locations, including protected areas, valleys, and hills, as well as watering sources, such as major and non-major water points and watering holes near rivers, which could vary in attractiveness and availability, and may further contribute to this discrepancy. This information, where available, can be incorporated into our model framework to improve its ability to predict movement and contact patterns more accurately.

The observed movement data was based on community participatory mapping, where local farmers identified resource areas on maps and reported movements [15]. This method is subject to recall bias, with the possibility of erroneous reports [27, 28]. We assumed that the loss of grazing and watering resources due to policy changes occurs randomly. While this approach provides some insights, it does not precisely capture the “land-squeeze” pattern resulting from socio-economic and political changes in Tanzania. In particular, changes in land use, fragmentation, and allocation of communal lands to conservation initiatives increasingly restrict areas available for grazing [29, 30]. Moreover, the rising livestock and human population indices in Tanzania in recent years may directly affect the herding capacity of agropastoralist livestock. The combined impact of these changes could potentially influence the predictions of our model.

Despite these limitations our models capture the essential features of current networks solely by using the spatial distribution of resources to generate simulated networks, a mechanism which reflects known drivers of agropastoralist decision-making. By projecting the outcomes of decisions under these same mechanisms under predicted future rainfall and vegetation conditions, we generate plausible future livestock movements patterns between villages. These patterns display both increased overall connectivity between villages due to climate-induced resource scarcity, but reductions in the importance of the most connected villages. As this both facilitates the spread of infectious diseases across the region and makes it harder to identify ‘important’ villages for disease control, reducing the impact of possible targeted interventions, it provides important considerations for the planning of future approaches to infectious disease prevention in a region which is likely to experience extreme challenges to animal and human health.

## 4 Methods

### 4.1 Data

This study uses three data sets: (1) geographical coordinates of villages and resource areas, together with the monthly observed network of intervillage connections through the use of shared resources; (2) NDVI data; and (3) climate data.

Location and observed network data were derived from a community participatory mapping study conducted by Ekwem et al. [15] between January and December 2017. The mapping exercise enabled the construction of monthly movement networks, representing village-to-village connections across shared resources such as grazing areas, water sources, dipping points, and salt points (Figure A2). The analyses in this current study focused on grazing and water, natural resources essential for livestock. We calculated Euclidean distance matrices to determine the distances between each village and its associated grazing and watering points.

Using a geographically weighted regression to model spatial variation in the relationship between historical NDVI, temperature, precipitation, and elevation, we predicted future trends in NDVI across the study area based on projected climate data (see supplementary materials). Historical NDVI data products were obtained from the third generation NOAA Global Inventory Monitoring and Modelling System (GIMMS3g), version 1 [31, 32]; and historical and projected climate data were obtained from the Climatologies at High Resolution for the Earth’s Land Surface Areas (CHELSA, https://chelsa-climate.org/).

### 4.2 Kernel movement model

We used a logistic function to model the probability that an individual livestock farmer within a village will select a specific resource location (grazing or watering) for their livestock. We selected this distribution for its simplicity and its ability to capture the decay in probability as distance increases. We included NDVI in our model equation to represent the attractiveness of resources and to capture the dynamic relationship between resource availability and herd movements. Consequently, the probability of movements from a village to a resource location is:

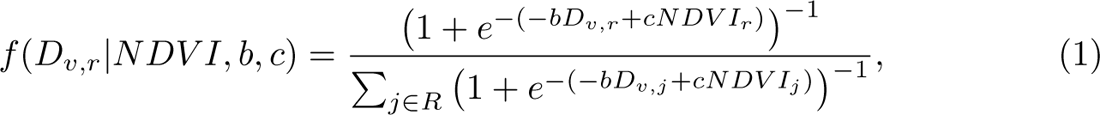

where *R* represents the set of all the locations of resources. The coefficients *b* and *c* indicate the strength of the distance and the effects of NDVI on the probability of selecting a resource location, respectively. As the value of *b* increases, the probability that a herd prefers a nearby location over a distant location also increases. The higher *c* is the more likely a herd is to prefer locations with high NDVI values, as they indicate more favourable grazing and watering conditions. To ensure that the movement probabilities form a valid probability distribution that sums to 1 for each source village, each probability to all possible destinations was divided by the sum of all probabilities.

#### 4.2.1 Monthly and seasonal movements

The observed movement networks presented in Figure A2 show varying patterns during different months, influenced by the availability of resources and changing environmental conditions. For example, in August and September, there is a substantial increase in contacts compared to any other month due to scarcity of resources. The persistence of common links between pairs of months in the grazing and watering networks can be seen in Figure A3. Months within the same season show higher proportions of repeated edges, indicating similar grazing and watering behaviours. To assess the significance of these repeated connections, we performed a randomisation test. This involved creating 1 000 monthly random networks based on the number of nodes and link density of the observed networks, using the *G*(*n, p*) algorithm of Erdős and Ŕenyi [33], where *n* is the number of nodes and *p* is the probability of a link between any two nodes. Then, we measured how many times the fraction of shared edges between any two months in the random networks is less than the observed networks. This yielded approximately 1, indicating that the observed monthly networks share significantly more common edges than expected by chance alone.

To incorporate these patterns in our model, we categorised movements into seasons and months: seasonal networks represent persistent links (which we shall simply refer to here as “static” links) within a season. In contrast, monthly networks represent dynamic links that change every month. The 12 months (*M*: January to December) were divided into four distinct seasons *S* based on rainfall patterns; wet (March, April, May, November, and December), short dry (February and July), transition (January, June, October), and long dry or drought (August and September) seasons. This categorisation facilitated the introduction of season-specific parameters *θ_s_* = (*b_s_, c_s_*) and month-specific parameters *θ_m_* = (*b_m_, c_m_*) for each season *s ∈ S* and month *m ∈ M*. Season-specific parameters allowed the generation of seasonal networks that reflect common structures shared among months within the same season, while month-specific parameters account for dynamic herd preferences and movements between different months. We assumed that herd movement patterns are consistent throughout any given month. Therefore, the probability that herds located in a village *v* select a resource location *r ∈ R* in a season or month *t ∈ {S, M}* is given by:

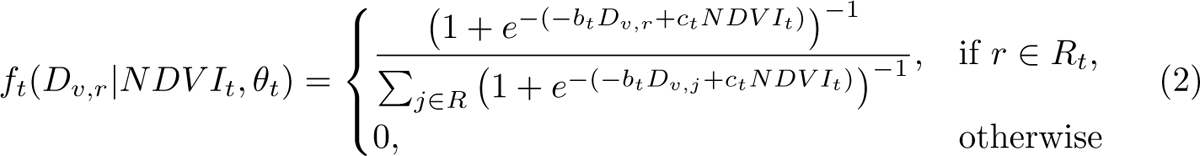

where *R* represents the set of all resource locations and *R_t_* = *{r ∈ R |* NDVI*_t_*(*r*) *>* 0.45*}* is the set of all resource locations available in a given season or month *t ∈ {S, M}*. Therefore, NDVI has the potential to affect grazing and watering movements through resource attractiveness and changing availability. Based on unpublished data, a loss of 15 *−* 20% in the number of grazing areas was observed during the long dry season (August and September) in 2016, which corresponds to a monthly NDVI threshold of 0.45.

### 4.3 Network generation

We used the estimated probability of movements in Equation 2 to simulate monthly and seasonal bipartite networks of village-to-resource locations. A bipartite network is a network whose nodes can be divided into two distinct sets *U* and *V* such that every link connects a node in *U* to one in *V* [34, 35]. These networks represent the connections between villages and resource points, where villages are the source nodes, resource points are the destination nodes, and movements from villages to resource points are the links. We used a multinomial distribution to stochastically assign network links from villages to specific resource points for each type of resource (grazing and watering) based on their estimated probability of movement.

In each village, multiple herds form groups and move to areas with resources. We assumed that the distribution of the number of monthly and seasonal herd movements across all villages follows a negative binomial distribution NB(*n_v_*; *µ_t_,* size*_t_*), with *µ_t_* as mean and size*_t_* as the dispersion parameter, where *n_v_* is the number of villages and *t ∈ {S, M}*. We chose the negative binomial distribution based on its flexibility in modelling overdispersion, which is important, especially during the wet season when there are limited movements to resource points. Each village *i* selects, with replacement, *n_i_ ∈* NB(*n_v_*; *µ_t_,* size*_t_*) resource locations from a multinomial distribution based on the probabilities derived from the parameterised Equation 2. As resources are selected with replacement, multiple herds from the same village can go to the same location, resulting in an adjacency matrix in which some values *≥* 1. In this simulation, the adjacency matrix *A^t^* is a rectangular *n_v_ × n_r_* matrix where the entry (*v, r*) indicates the number of herds from village *v* that use the resource point *r*, and zero when no herd from village *v* uses the resource point *r*. Since our focus is on contacts between villages at shared resources in a given month or season rather than the precise number of contacts, when more than one herd from a village moves to the same location, the corresponding entry in the adjacency matrix *A^t^* takes the value 1. This process transforms the adjacency matrix into a binary representation that shows the presence or absence of a link between a village and a resource point in a given month or season. In the matrix to the left of Equation 3, the elements *p_ij_* represent the probabilities of movement from a village *i* to the resource point *j*:

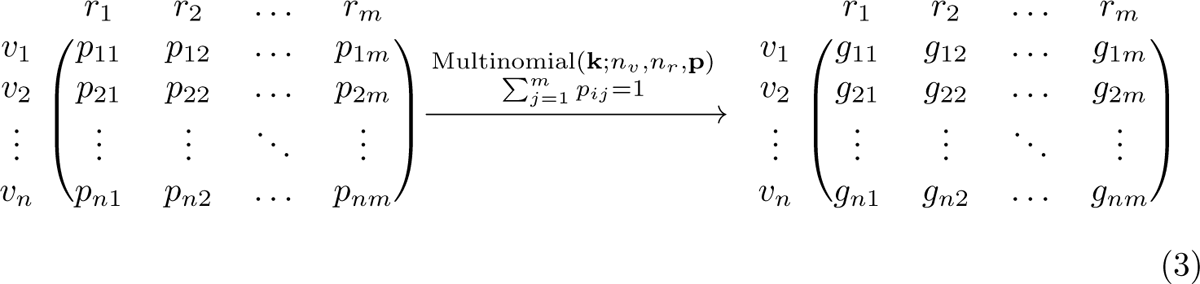

After applying the multinomial distribution, the second matrix in Equation 3 is obtained, where *g_ij_* represents the links of the herds that travel from the village *i* to the resource point *j*. We simulated monthly and seasonal networks for each resource type and aggregated them (using the network union operation) to capture static and dynamic network links.

### 4.4 Approximate Bayesian computation

We used approximate Bayesian computation (ABC) based on a rejection sampling algorithm [36, 37], as implemented in the “EasyABC” package [38] in R [39], to estimate model parameters (Table A1). We conducted 200 000 iterations of the ABC algorithm for each resource type, month, and season. In each iteration, we sampled parameters from the prior distributions listed in Table A3, ran the model three times to adjust for stochastic variation, and calculated the mean of eight summary statistics. These included the number of active nodes (defined as nodes that are connected to at least one other node in the network), mean degree centrality, mean betweenness centrality, mean eigenvector centrality, assortativity index, mean geographic distance between connected villages, and the number of links formed between villages less than 8 km and less than 15 km apart. We chose these summary statistics to capture the key characteristics of the observed village-village networks. The summary statistics of the simulated networks were normalised using the standard deviation of the summary statistics across previous model simulations before calculating the Euclidean distance between the observed and simulated data [38]. The tolerance, which determines the proportion of parameters kept, was set to 0.001, resulting in a posterior distribution with 200 parameters.

### 4.5 Posterior predictive distribution and network analysis

To evaluate the performance of our network simulation model, we compared the observed networks and the simulated bipartite networks for each resource type at both the monthly and the yearly levels. We randomly drew parameters from the posterior distributions to parameterise the model and generated 1 000 monthly bipartite networks for each resource type, of which 100 were sampled for further analysis. We selected 100 simulations since it was observed that most centrality measures became stable after this number of simulated networks (see Figures A6 and A7). We then transformed each simulated bipartite network into a one-mode village-to-village network representation.

We used a variety of network measures, both at the node level and at the network level, to evaluate the structural properties of these networks. At the node level, we calculated centrality measures, including degree centrality, betweenness centrality, and eigenvector centrality. In addition, we calculated network-level measures such as node count, transitivity (clustering coefficient), mean distance, network diameter, and degree assortativity index (see description in Table A2).

### 4.6 Generating future NDVI

We used a geographically weighted regression (GWR) to model spatial variations in the relationship between NDVI and climate variables (temperature and precipitation) in the Serengeti district. The coefficients of the GWR model were combined with CMIP6 climate projections to estimate the future monthly values of NDVI in the Serengeti district (see Appendix B for additional information).

### 4.7 Generating future networks

We explored the long-term impacts of future climate change scenarios on network structures by incorporating projected NDVI values into a parameterised network generation model. This allowed us to simulate future network scenarios, using 2050 and 2080 as examples. As the pattern of the impact of policy changes on the loss of grazing and watering locations remains unclear, we simulated a scenario where resources were lost randomly, with a 10% reduction as an illustration.

## Appendix A Supplementary methods

### A.1 Study area

This analysis focused on the Serengeti district in northern Tanzania (Figure A1), with an estimated human population of 249 420 [40]. Households in the district practise agropastoralism [41], which combines crop farming and livestock-keeping [2], with the primary sources of income being crops, supported by income from livestock-related sales [1]. Agropastoralists in this area engage in daily movements of livestock (cattle, sheep, and goats) to access shared resources at different locations [5].

**Table A1:**
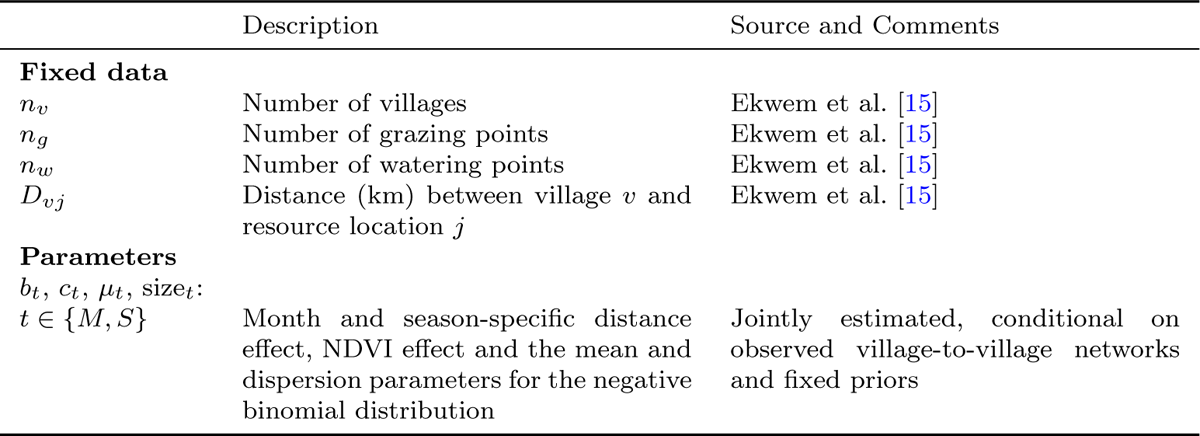
Summary of model data and fixed parameters. We used a Bayesian framework to estimate month- and season-specific parameters *θ_m_* and *θ_s_* for each season and month, which provides a parameter estimation approach that accounts for and quantifies the parameter uncertainty in model predictions by producing parameter distributions rather than point estimates.

## Appendix B Climate Change Impacts on Vegetation Dynamics in Serengeti District

### B.1 Introduction

The changing climate poses a significant and complex challenge for the vegetation in regions like northern Tanzania [25, 29, 30]. This challenge has potential consequences not only for local ecosystems but also for the livelihoods of livestock farmers who depend on these natural resources [1, 7, 10, 29, 50]. The impacts of climate change on vegetation in this part of the world and other sub-Saharan countries are primarily driven by rising temperatures, altered rainfall patterns, and increasing climate variability [25]. These factors directly influence the growth and distribution of vegetation and water availability, affecting the well-being of livestock herds [6, 25]. However, with ongoing changes in climate conditions, how vegetation will respond remains a critical question, and there is a scarcity of detailed studies that project the future of vegetation in this highly vulnerable region.

**Fig. A1:**
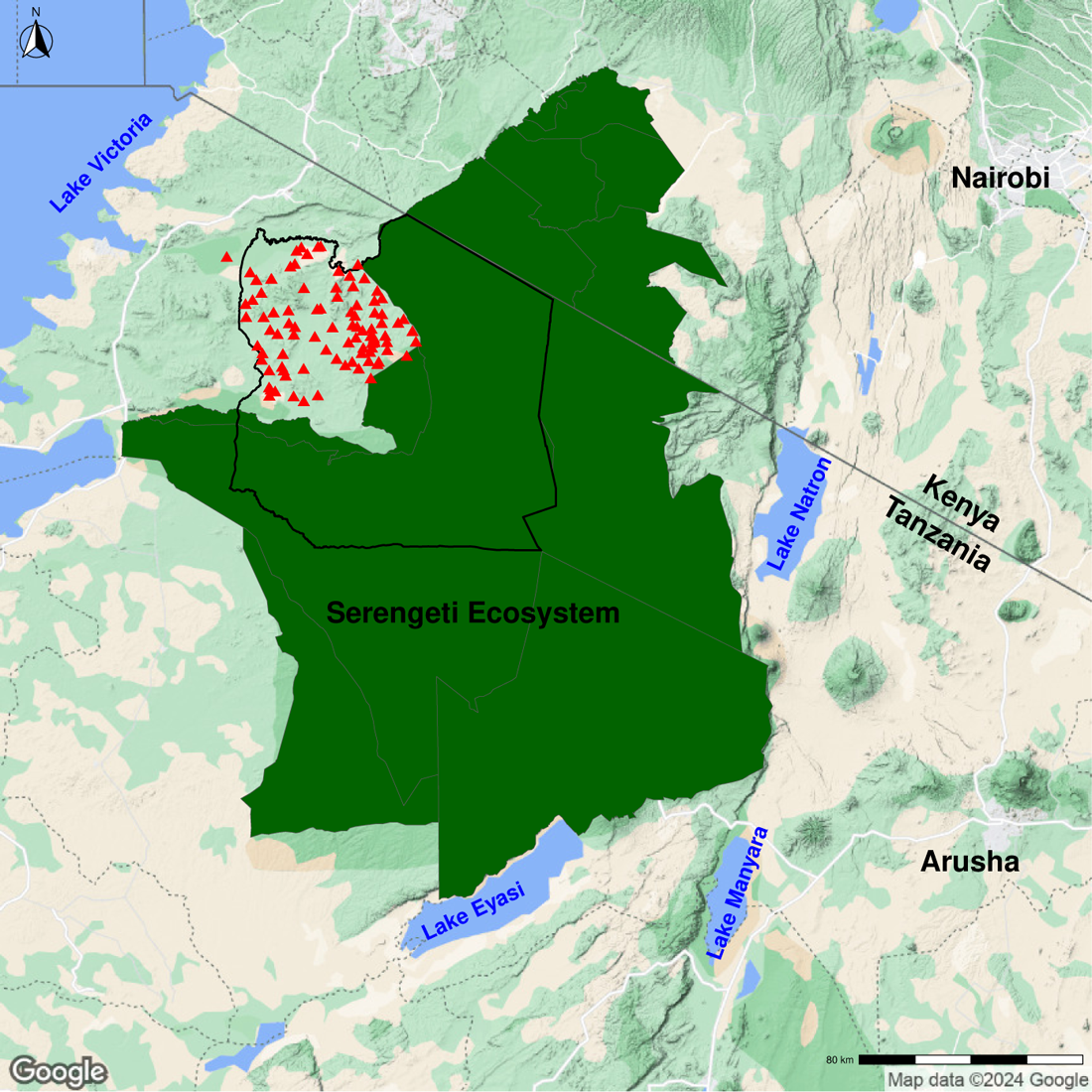
Map of the study area in northern Tanzania showing key features. The protected areas of the Serengeti ecosystem are shaded in dark green, sourced from shapefiles available at https://serengetidata.weebly.com. The black line marks the boundary of the Serengeti region, and the red triangular points show village locations inside the region [41]. The map’s background is a Google Static Terrain map, accessed using the *ggmap* package in R [42]. This map was created in R [39] using the *ggplot2* and *terra* packages [43, 44].

Recently, there has been an increase in interest in understanding the interactions between vegetation and climate change. Climate data can be integrated with vegetation data in a statistical model to predict future changes in vegetation dynamics. This approach has shown promise in predicting future leaf area index (LAI) trends [51, 52], NDVI trends [53], and short-term prediction of NDVI as an early warning system for disasters such as famine and epidemic diseases [54], and the risk of Rift Valley fever (RVF) [55, 56]. These methods help our understanding of ecological dynamics and facilitate informed decision-making for sustainable resource management.

**Fig. A2:**
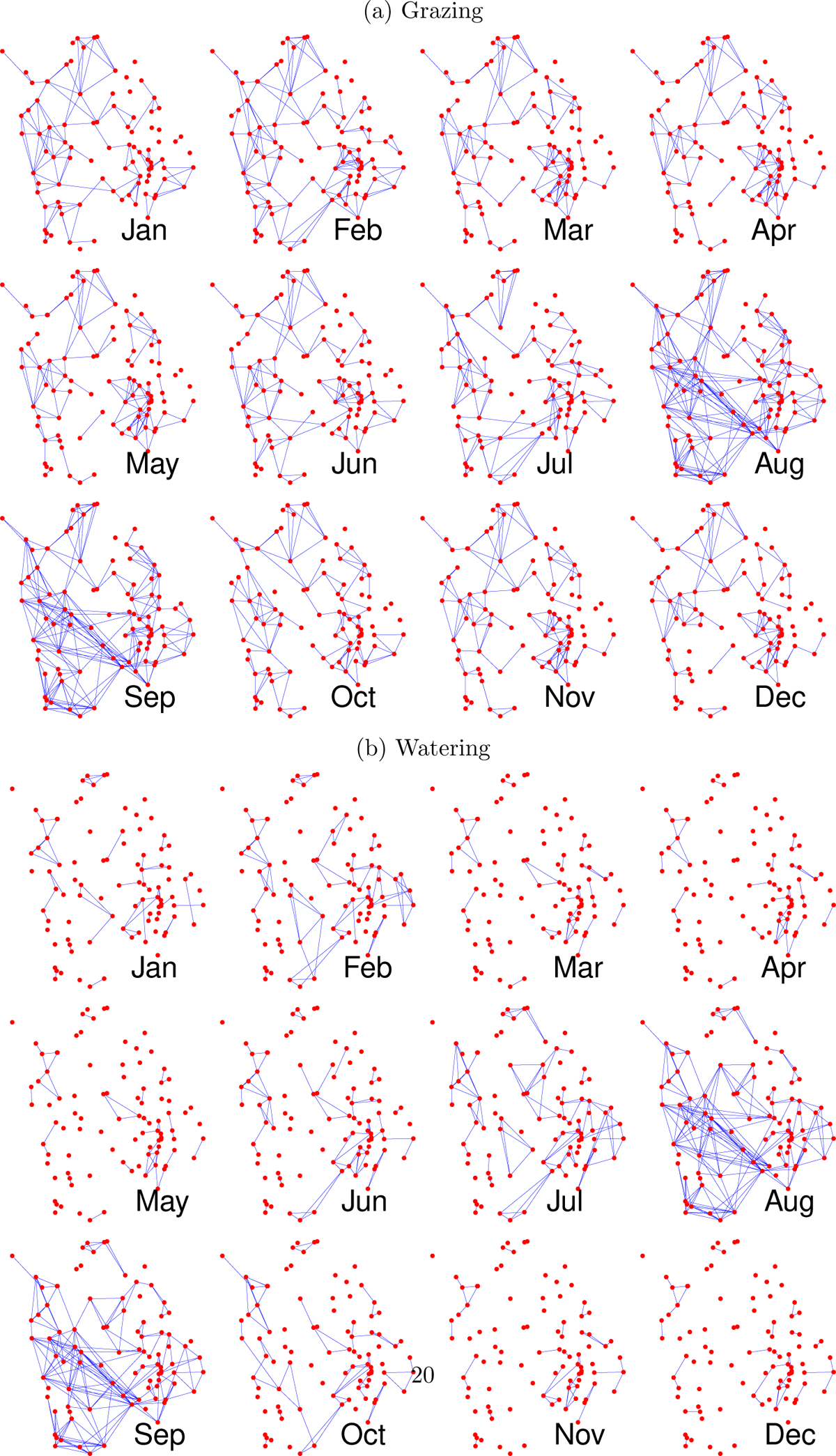
Observed spatial livestock movement networks of village connectivity showing variation in contact patterns across months and shared resource type [(a) grazing and (b) watering)] in Serengeti district, northern Tanzania. Nodes (red points) are villages in their geographical position, and the blue edges represent connected villages using shared resource areas.

**Fig. A3:**
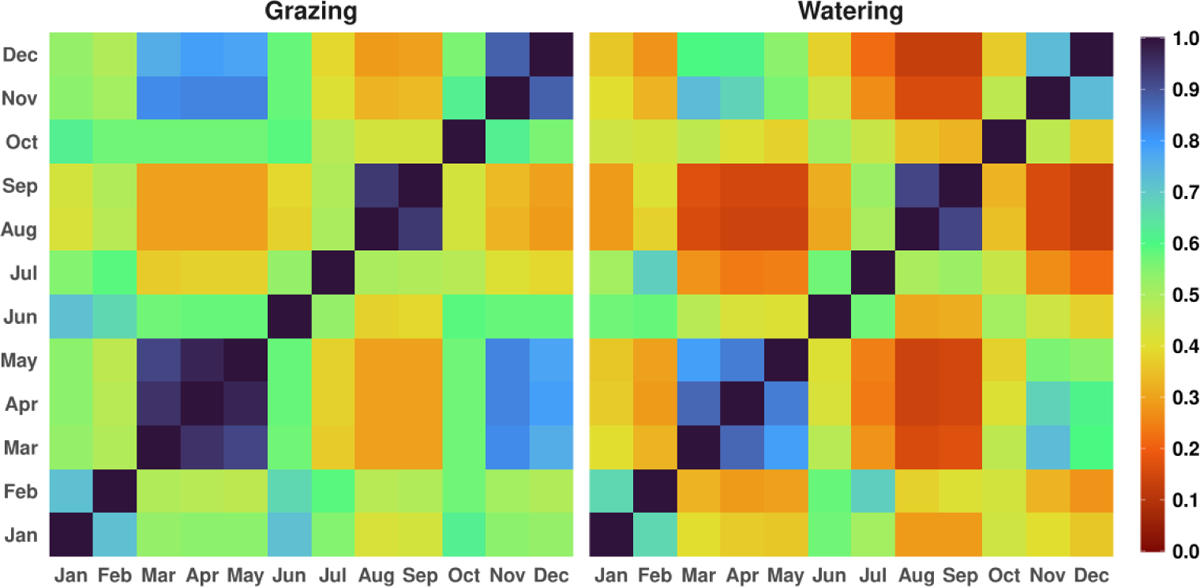
Variation in the relative proportion of common links shared between each pair of observed monthly grazing networks (left) and watering networks (right). Each entry *ij*, where *i, j ∈ {*Jan, …, Dec*}*, is the number of common links in the row network *G_i_* and column network *G_j_*, divided by the total number of unique links in both networks, i.e., 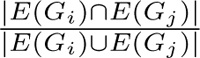.

The primary purpose of this analysis is to investigate the relationship between climate factors and vegetation and project how vegetation might change in the future. We used the data generated here in the parameterised movement model (4.7) to simulate future network scenarios. Our approach uses a geographically weighted regression (GWR), which uses historical climate data, consisting of temperature and precipitation, to predict future trends in the NDVI within the Serengeti district in the Mara region of Tanzania. This method captures spatial heterogeneity in climate-vegetation relationships by allowing the relationships between independent and dependent variables to vary locally. We predicted future vegetation in the Serengeti region based on the GWR model and future climate projections from a model in the Coupled Model Intercomparison Project Phase 6 (CMIP6) under a high-emission scenario (SP5850).

### B.2 Method

#### B.2.1 Study area

The study area for this analysis was the Serengeti district in northern Tanzania (Figure B8a). Monthly historical temperature, precipitation, and NDVI across the study area from 1982 to 2014, with the overall monthly mean, are presented in Figure B9 and Figure B8b. Low precipitation values are observed from June to October, corresponding to low mean NDVI values in the following months, demonstrating the impact of rainfall on vegetation growth. Most of this area is rural, and most households depend on both crop farming and livestock keeping (agropastoralism) for their livelihoods [5]. The Serengeti ecosystem plays a crucial role in providing grazing lands for livestock.

**Table A2:** Description of network measures.

| Network Measure | Description |
| --- | --- |
| Degree centrality | Number of connections each village has [26] |
| Betweenness centrality | Measures the extent to which a village lies on the shortest path connecting other villages of the network [45] |
| Eigenvector centrality | Assigns importance scores to villages based on both their centrality and the centrality of their neighbouring villages [46]. Unlike degree centrality, which treats every contact equally, it treats contacts according to their centralities, so that connecting to some villages has more value than connecting to others [47] |
| Node count | Represents the size of the network [26] |
| Transitivity | Indicates the tendency of a village to form tightly connected clusters [48] |
| Mean distance | Mean pairwise shortest network distance between the villages of the network [26] |
| Network diameter | Represents the length of the longest shortest path (network distance) between two villages [26] |
| Degree assortativity | Indicates the tendency of villages to connect with others of similar degrees [49] |

**Table A3:** Uniform prior distribution of parameters across different months and seasons. Values within parentheses represent the range of uniform priors.

| Month | Grazing |  |  |  | Watering |  |  |  |
| --- | --- | --- | --- | --- | --- | --- | --- | --- |
|  | b | c | Shape | Mean | b | c | Shape | Mean |
| Jan | (0.6, 8) | (0, 1) | (0.01, 5) | (0.1, 3) | (0.6, 8) | (0, 1) | (0.01, 5) | (0.1, 3) |
| Feb | (0.5, 8) | (0, 1) | (0.01, 5) | (0.1, 4) | (0.5, 8) | (0, 1) | (0.01, 5) | (0.1, 4) |
| Mar | (0.8, 8) | (0, 1) | (0.01, 5) | (0.1, 2) | (0.8, 8) | (0, 1) | (0.01, 5) | (0.1, 2) |
| Apr | (0.8, 8) | (0, 1) | (0.01, 5) | (0.1, 2) | (0.8, 8) | (0, 1) | (0.01, 5) | (0.1, 2) |
| May | (0.8, 8) | (0, 1) | (0.01, 5) | (0.1, 2) | (0.8, 8) | (0, 1) | (0.01, 5) | (0.1, 2) |
| Jun | (0.6, 8) | (0, 1) | (0.01, 5) | (0.1, 3) | (0.6, 8) | (0, 1) | (0.01, 5) | (0.1, 3) |
| Jul | (0.5, 8) | (0, 1) | (0.01, 5) | (0.1, 4) | (0.5, 8) | (0, 1) | (0.01, 5) | (0.1, 4) |
| Aug | (0.3, 8) | (0, 1) | (0.01, 5) | (0.1, 1) | (0.3, 8) | (0, 1) | (0.01, 5) | (0.1, 1) |
| Sep | (0.3, 8) | (0, 1) | (0.01, 5) | (0.1, 0.5) | (0.3, 8) | (0, 1) | (0.01, 5) | (0.1, 0.5) |
| Oct | (0.6, 8) | (0, 1) | (0.01, 5) | (0.1, 4) | (0.6, 8) | (0, 1) | (0.01, 5) | (0.1, 3) |
| Nov | (0.8, 8) | (0, 1) | (0.01, 5) | (0.1, 2) | (0.8, 8) | (0, 1) | (0.01, 5) | (0.1, 2) |
| Dec | (0.8, 8) | (0, 1) | (0.01, 5) | (0.1, 1) | (0.8, 8) | (0, 1) | (0.01, 5) | (0.1, 1) |
| Season |  |  |  |  |  |  |  |  |
| Wet | (0.8, 8) | (0, 1) | (0.01, 5) | (0.5, 4) | (0.8, 8) | (0, 1) | (0.01, 5) | (0.5, 5.5) |
| Dry | (0.5, 8) | (0, 1) | (0.01, 5) | (0.5, 6) | (0.5, 8) | (0, 1) | (0.01, 5) | (0.5, 7.5) |
| Transition | (0.6, 8) | (0, 1) | (0.01, 5) | (0.5, 6) | (0.5, 8) | (0, 1) | (0.01, 5) | (0.5, 7) |
| Drought | (0.3, 8) | (0, 1) | (0.01, 5) | (1.0, 6) | (0.3, 8) | (0, 1) | (0.01, 5) | (1.0, 8) |

**Fig. A4:**
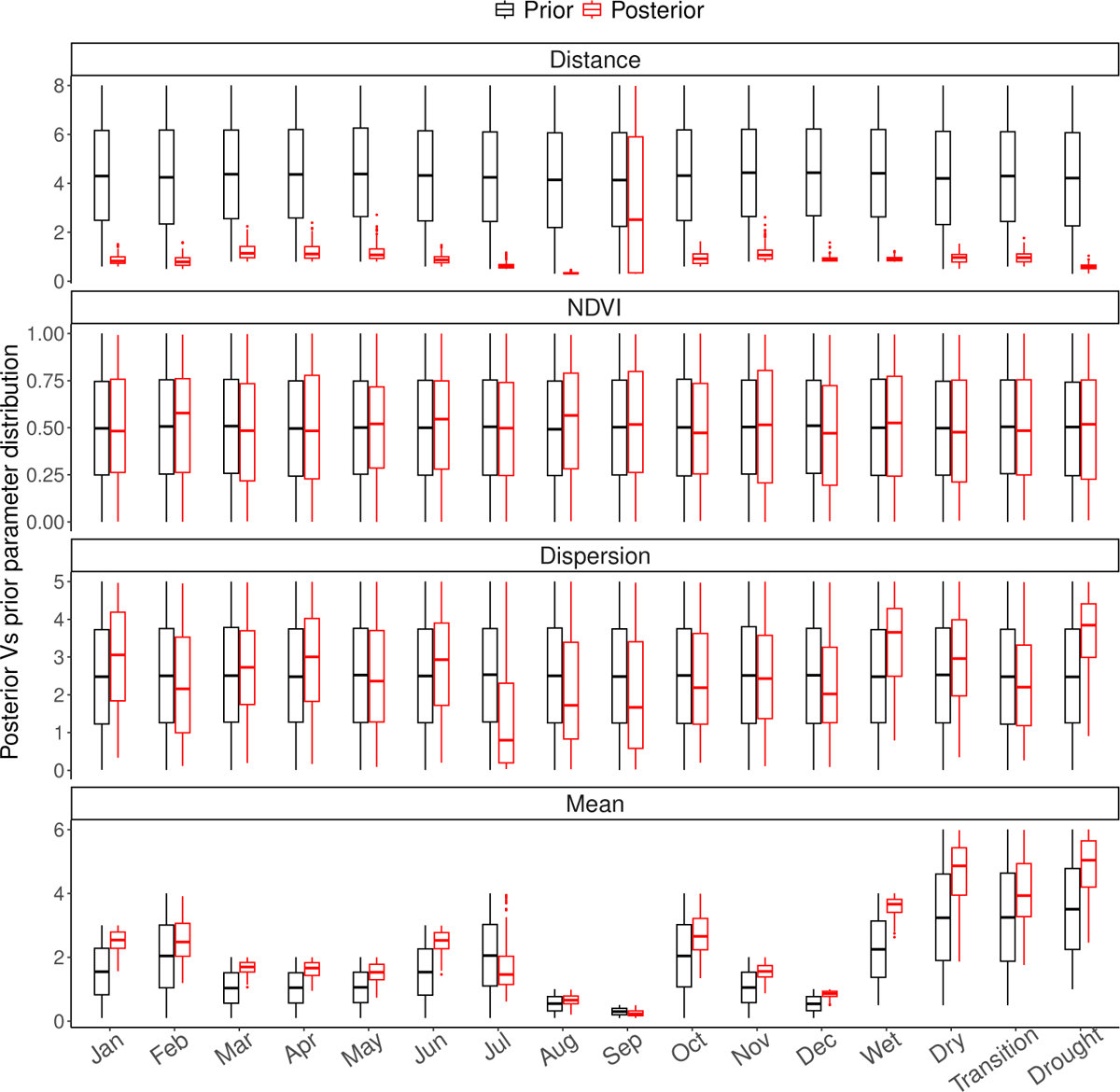
The posterior distributions (red) of grazing model parameters by month and season estimated by fitting the kernel movement model to summary statistics derived from observed movement network data, using ABC and priors (black). The median values of the parameter estimates are represented by the red central dashed lines in the figure.

#### B.2.2 Data

We obtained the NDVI data product of the third-generation NOAA Global Inventory Monitoring and Modelling System (GIMMS3g), version 1 [31]. GIMMS data have a spatial resolution of 1*/*12 degrees (*≈* 10 km) and a bimonthly temporal resolution, covering from July 1981 to December 2015. It is widely used to analyse long-term vegetation changes in various regions [31] and is publicly available [32]. Here, we focus our analysis on monthly NDVI data from 1982 to 2014. Elevation data were acquired from the Shuttle Radar Topography Mission global 3 arc-second (SRTM GL3) available through the “Elevatr” package in R, with a spatial resolution of 90 m [57].

Historical climate data on monthly temperature and precipitation from 1982 to 2014 were obtained from the Climatologies at High Resolution for the Earth’s Land Surface Areas (CHELSA, https://chelsa-climate.org/, accessed on 30 September 2023). This data set has a spatial resolution of 1*/*120 degrees (*≈* 1 km) and is based on a statistical downscaling algorithm of the global reanalysis data [58]. Finally, monthly temperature and precipitation projections with a spatial resolution of 1 km were extracted using the CHELSA algorithm [58, 59]. These projections were based on simulations from the Met Office Hadley Centre (MOHC) HadGEM3-GC31-MM model under high emission scenarios (SSP585) as part of the sixth phase of the Coupled Model Intercomparison Project (CMIP6, https://esgf-index1.ceda.ac.uk/search/cmip6-ceda/). The projections cover the period from 2015 to 2100. All datasets were resampled to a resolution of 0.025 (*≈* 2.775 km) degrees using bilinear interpolation.

**Fig. A5:**
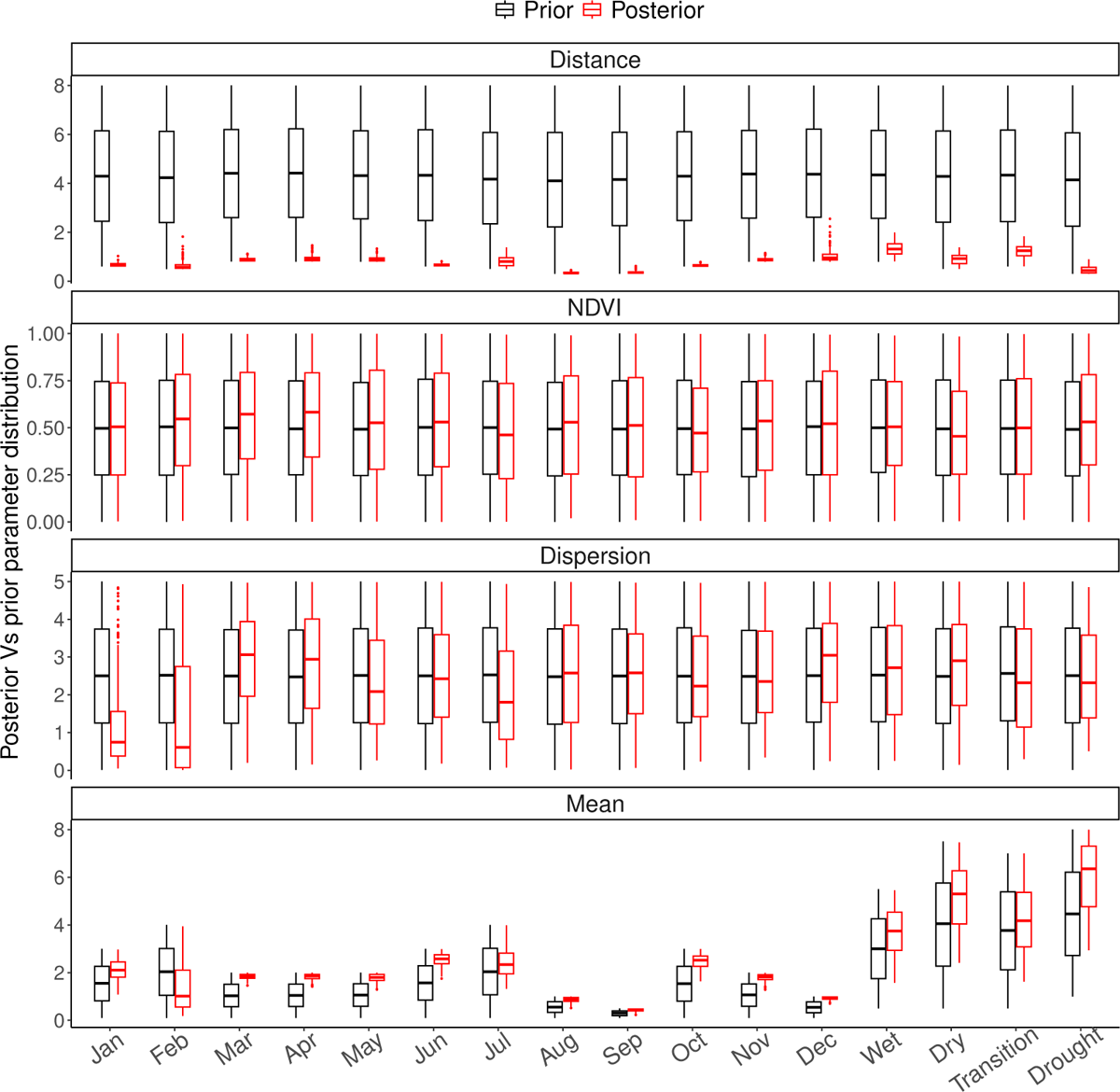
The posterior distributions (red) of watering model parameters by month and season estimated by fitting the kernel movement model to summary statistics derived from observed movement network data, using ABC and priors (black). The median values of the parameter estimates are represented by the red central dashed lines in the figure.

**Fig. A6:**
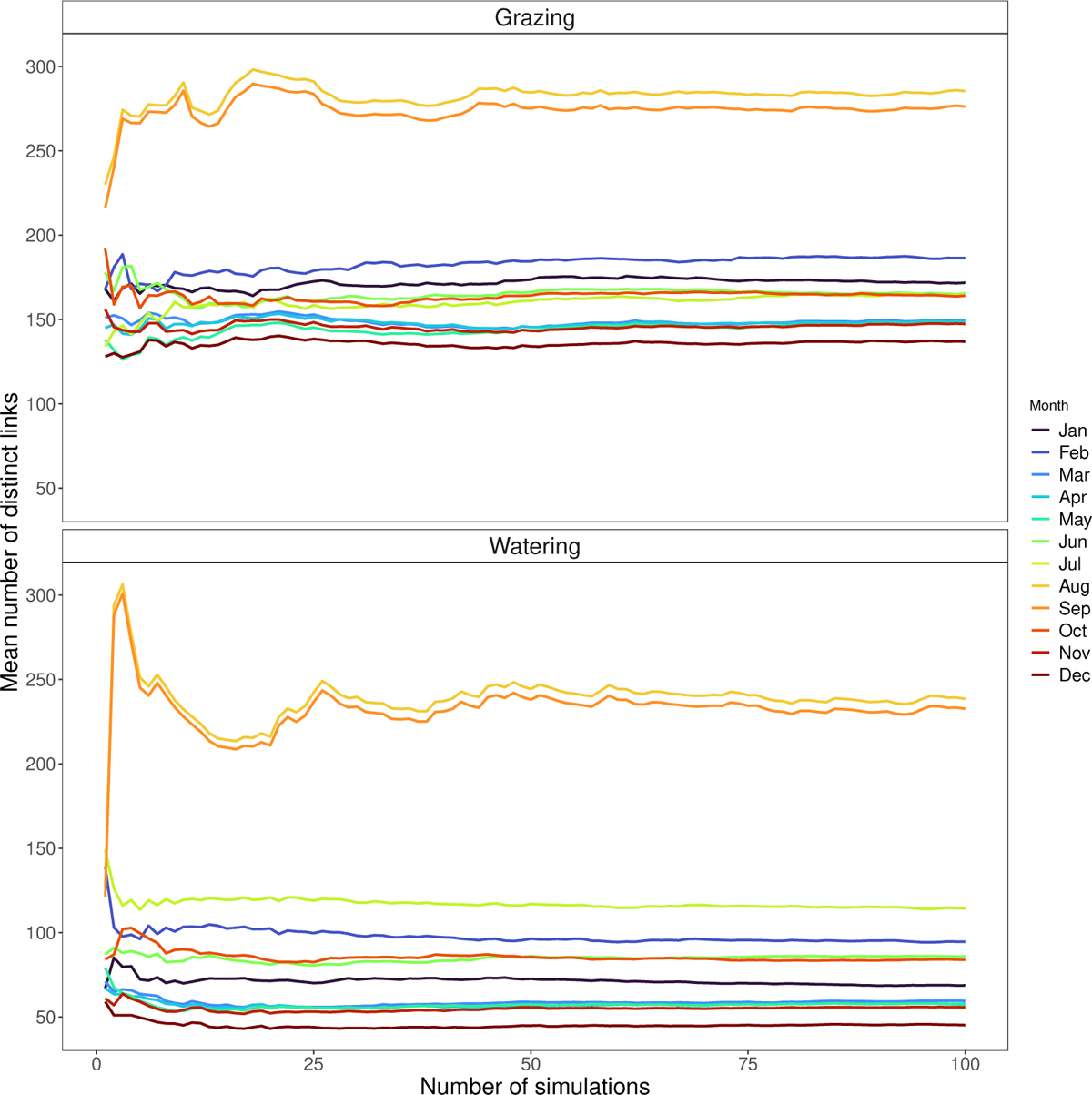
Convergence of the monthly number of unique links after 100 simulations.

**Fig. A7:**
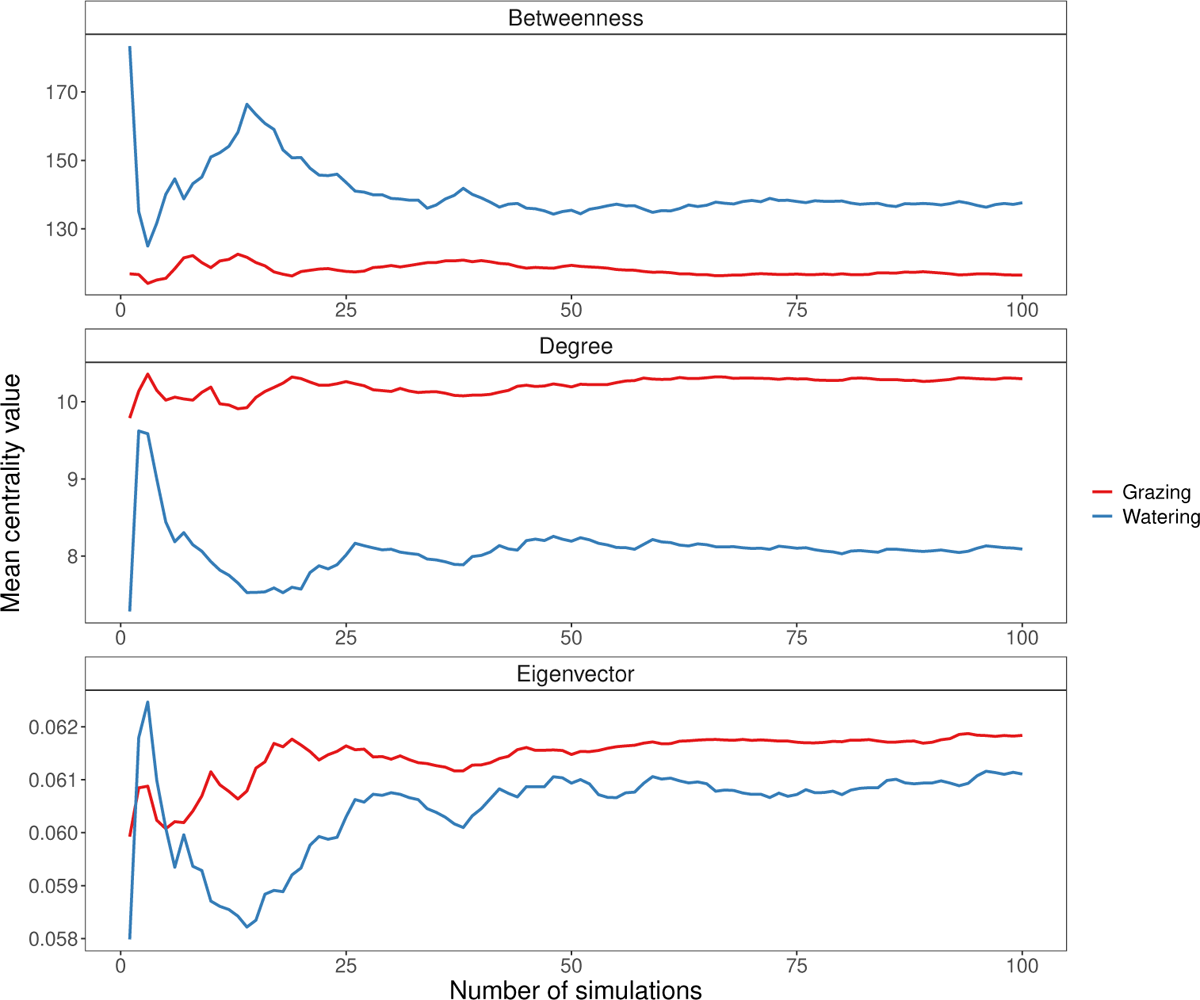
Convergence of mean degree, betweenness, and eigenvector centralities in year grazing and watering networks after 100 simulations.

#### B.2.3 Regression Model

We used multiple linear regression to determine the relationship between NDVI, climate factors (temperature and precipitation), and elevation. The equation for this is:

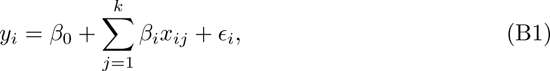

where *y_i_* is the dependent variable, *x_ij_* is the value of the *j^th^* independent variable, *k* is the number of independent variables, *β*_0_ is the intercept, *β_i_*is the slope coefficient for the *k^th^* independent, and *ɛ* is the random error that is normally distributed with zero mean and common variance *σ*^2^. Here, the dependent variable is NDVI, while the independent variables are temperature, precipitation, elevation, and month.

We fitted an Ordinary Least Squares (OLS) regression across the study area. We used monthly historical data spanning 33 years from 1982 to 2014, including temperature, precipitation, elevation, and sinusoidal representations of months as independent variables. The global regression model had an adjusted *R*^2^ value of 0.5374, indicating that the independent variables could explain a considerable amount of the NDVI variance in the region. However, the model also revealed significant spatial clustering of residuals.

Previous research has shown that vegetation responses to climate variations differ between locations and vegetation types [60–64]. A single global regression model assumes a stationary relationship between NDVI and climate variables across the study area. This may lead to bias and an inaccurate representation of local dynamics [60, 65]. We used a geographically weighted regression (GWR) [65] to address potential spatial variation in the dependent-independent variable relationships. This allows separate local regressions for each location (grid cell) in the study area. Each local regression uses the dependent and explanatory variables from the neighbourhood surrounding the target location [65, 66]. Bandwidth determines the size of the neighbourhood and can be calculated using statistical methods such as cross-validation (CV) or the Corrected Akaike Information Criterion (AICc) [65].

We used AICc to determine the optimal bandwidth. AICc is preferred to CV because it reflects model parsimony and prevents overfitting of GWR models [67]. The bandwidth, which defines the proportion of observations included in each local model, adapts based on the spatial distribution of the observations. This means that the bandwidth adjusts according to the density of observations, ensuring that each local estimate relies on a consistent number of nearby data points [65, 67]. We used a bisquare kernel to calculate distance weights, giving higher weights to nearby observations to compute local model coefficients. The GWR model produces spatially varying model coefficients, and the performance of each local model is evaluated by a local *R*^2^. The overall performance of the GWR model is assessed by the overall *R*^2^. We used the “GWmodel” package [66, 67] to perform the GWR analysis in R version 4.3.1 [39].

**Fig. B8:**
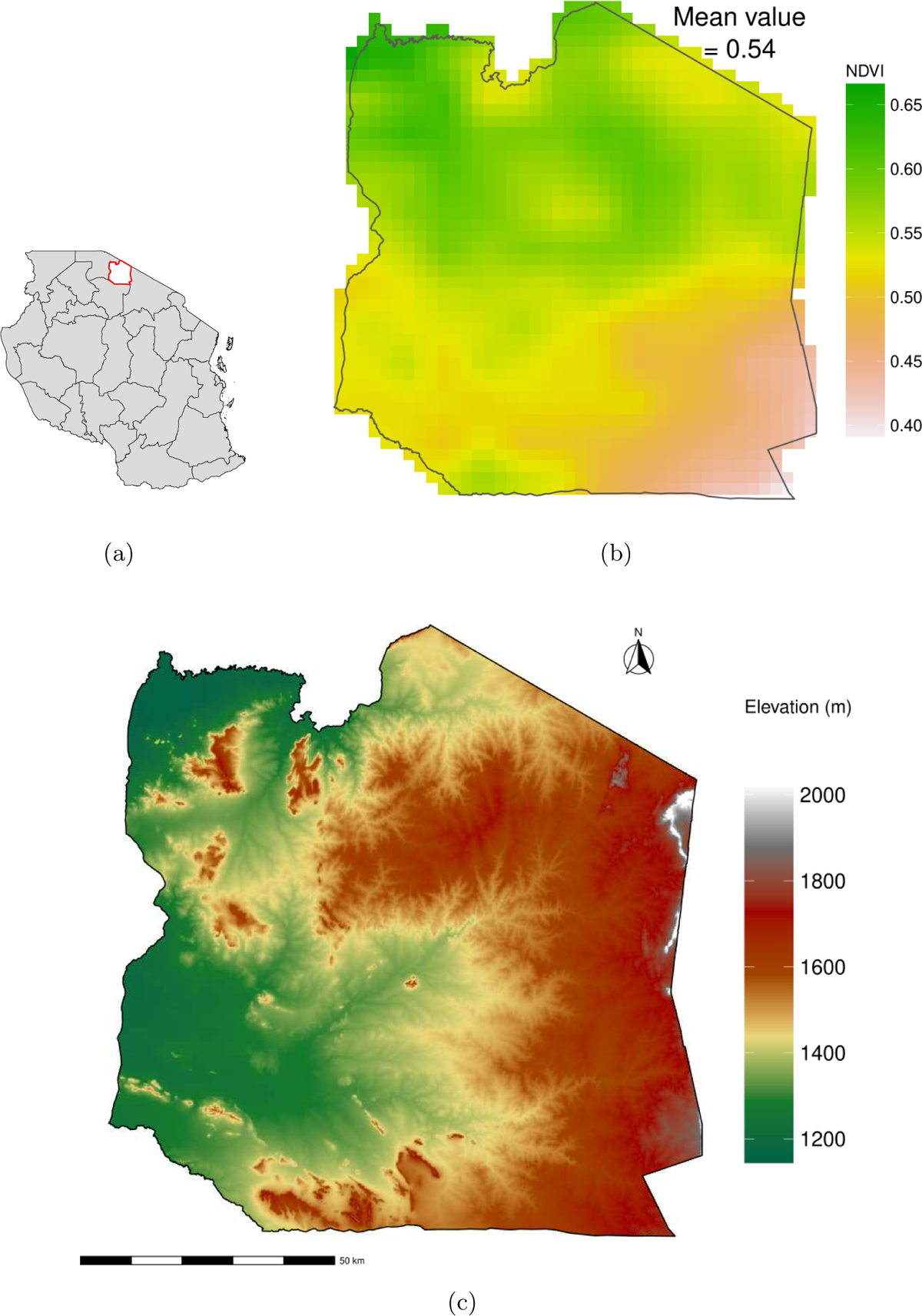
Tanzania map with a red boundary representing the study area (a), elevation (b), and spatial mean historical NDVI from 1982 to 2014 (d). The boundary shape file was downloaded from https://gadm.org/ using the “geodata” package available in R.

**Fig. B9:**
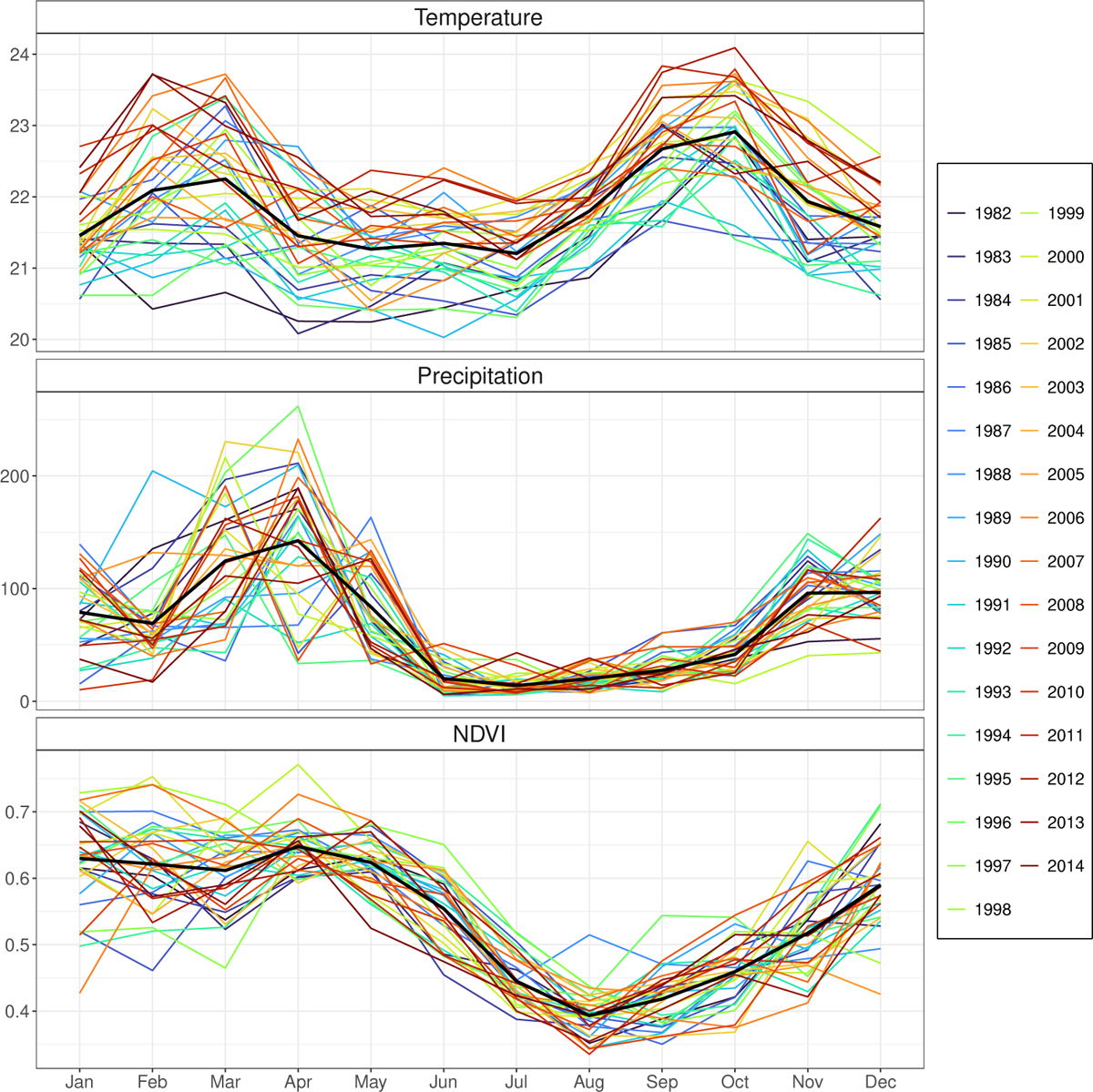
Yearly temperature, precipitation, and NDVI from 1982 to 2014 for each month and their corresponding monthly mean across all years shown as a black line.

### B.3 Results

#### B.3.1 GWR model

The GWR model results are presented in Figure B10, including the spatial distribution of residuals (Figure B10a), local *R*^2^ Figure B10b, intercepts (Figure B10c), precipitation (Figure B10d), temperature (Figure B10e), and elevation (Figure B10f). The GWR model substantially improved our ability to capture spatial variations in NDVI, achieving an overall *R*^2^ value of 0.6775. The spatial distribution of the local *R*^2^ showed the effectiveness of the local regression models in fitting the observations (Figure B10b). On average, within each area determined by bandwidth (1.6%), temperature, precipitation, and elevation could explain *≈* 68% of the variation in NDVI.

Approximately 74% of the local *R*^2^ values were greater than 0.6, particularly in the eastern region with high elevation. In contrast, the western region mainly consists of grazing and cropping activities with lower local *R*^2^ values. The temperature coefficients ranged from *−*0.023 to 0.012, with 90% of the region having negative coefficients from the GWR model. However, the precipitation coefficients were mainly positive, ranging from *−*0.01 to 0.02, with 92% of the region having positive values. Elevation coefficients ranged from *−*0.0014 to 0.0014, with most regions having negative values.

#### B.3.2 Future NDVI predicted by GWR model

Historical and projected mean temperature, precipitation, and NDVI trends from 1982 to 2100 are shown in Figure B11. Under the SSP585 scenario, the Serengeti district is expected to experience an increase in mean temperature and precipitation. Notably, there is a consistent upward temperature trend from the reference period (1982–2014) to the late century. During the reference period, the mean temperature was 21.83*^◦^*C, ranging from 17.06*^◦^*C to 24.41*^◦^*C (Table B4). As we move into the early century (2015– 2040), the mean temperature rises to 22.86*^◦^*C (with a range of 18.22*^◦^*C to 25.44*^◦^*C). By mid-century (2041–2070), the mean temperature increased further to 24.66*^◦^*C (with a range of 19.35*^◦^*C to 27.61*^◦^*C). In the late century (2071–2100), the mean temperature is projected to increase substantially to 27.23*^◦^* C (with a range of 21.59*^◦^*C to 30.58*^◦^*C), indicating a substantial warming trend.

**Fig. B10:**
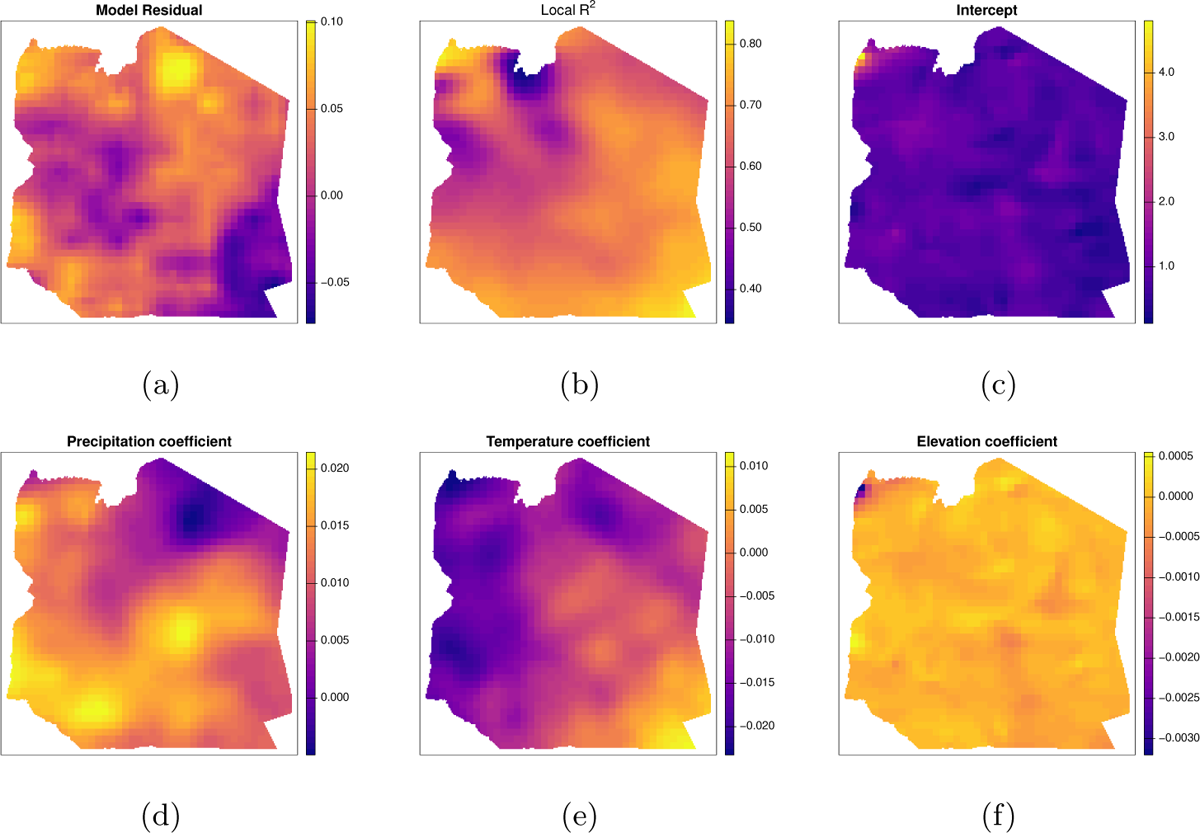
Spatial distribution of model residuals (a), local *R*^2^ (b), intercept (c), and model coefficients for precipitation (d), temperature (e) and elevation (f).

While mean precipitation values exhibit some fluctuations, a noticeable increase in precipitation levels emerges as we transition to the late century. The mean monthly precipitation was 67.92 mm during the reference period, ranging from 32.15 to 136.51 mm. In the early century, this value increased to 72.88 mm (with a range of 36.62 mm to 148.62 mm). By the mid-century, the mean precipitation increased to 81.23 mm (with a range of 40.29 mm to 164.26 mm) and increased further to 93.10 mm (with a range of 48.93 mm to 195.28 mm) in the late century. These values represent an approximate increase of 7.3%, 19.6%, and 37.1%, respectively, compared to the reference period.

**Table B4:**
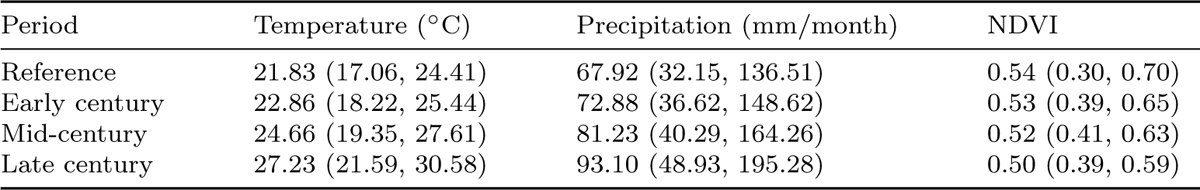
Historical (1982 – 2014) and future (2015 – 2100) mean values and spatial ranges for yearly temperature, precipitation, and NDVI in Serengeti.

NDVI values exhibit a declining trend over the studied time frames. In the reference period, the mean NDVI was 0.54 (with a range of 0.3 to 0.7). By the early century, the mean NDVI decreased to 0.53 (with a range of 0.39 to 0.65), indicating an approximate 1.9% decrease compared to the reference period. Moving into the mid-century, the mean NDVI declines further to 0.52 (with a range of 0.41 to 0.63), marking an approximate 3.7% decrease. By the late century, the mean NDVI is projected to decrease to 0.50 (with a range of 0.39 to 0.59), representing an approximate 7.4% decrease in the mean NDVI over the century.

The Figure B13 illustrates the spatial distribution of future changes in NDVI relative to the baseline period (1982–2014). The changes in NDVI are more pronounced in the late century than in the mid-century. The western and northern parts of the Serengeti are expected to experience a substantial decrease in NDVI, while the southeastern region is likely to experience an increase in NDVI. This suggests that vegetation dynamics will be more variable in the future, with increases and decreases more prominent in the late century.

**Fig. B11:**
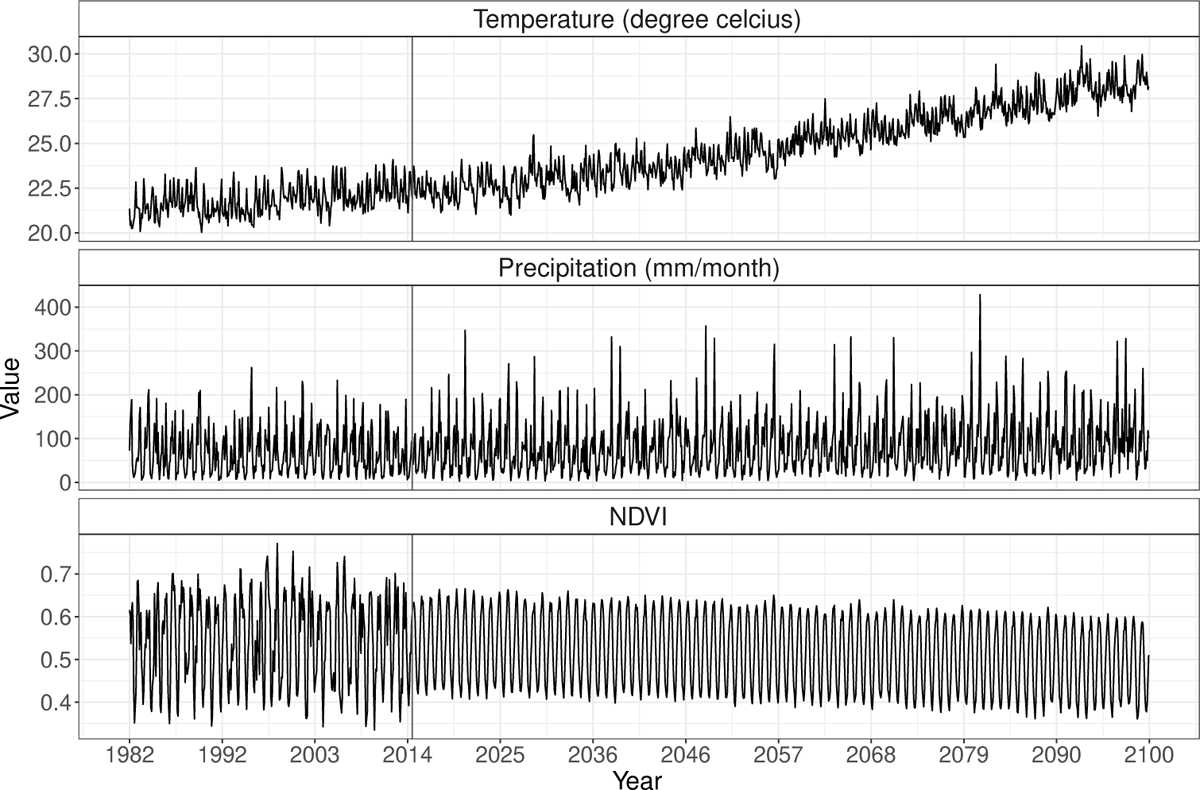
Time series of monthly mean temperature, precipitation, and NDVI across study area for historical (1982 – 2014) and future (2015 – 2100) under the SSP585 scenario. The blue and red lines are trend lines of historical and projected NDVI values. The vertical grey line divides the time series into historical and future.

**Fig. B12:**
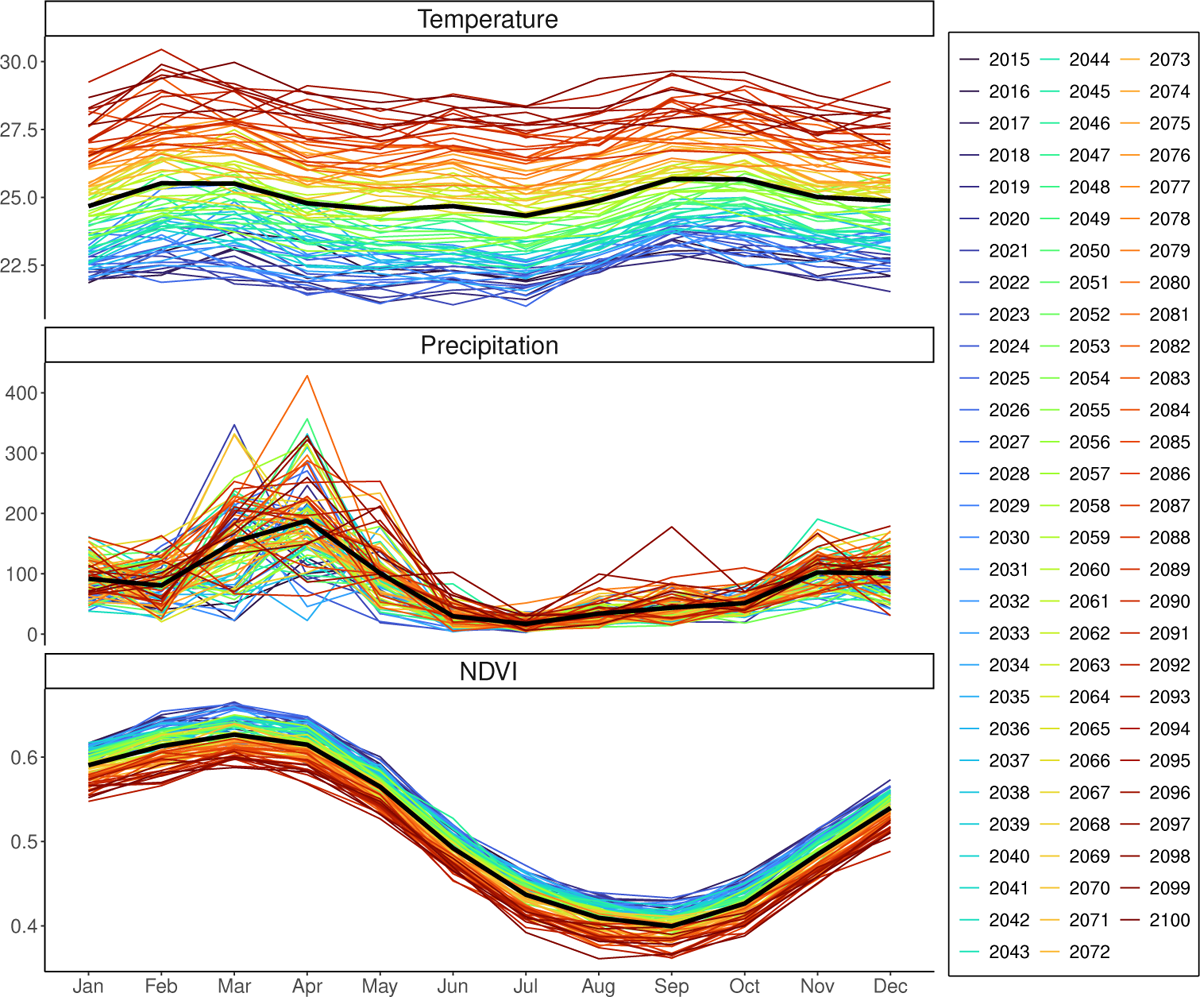
Monthly variation in temperature, precipitation, and NDVI for each year from 2015 to 2100. Each coloured line represents the mean value over all grid cells for each year, while the black line represents the mean value across all years from 2015 to 2100.

### B.4 Discussion

In this study, we used geographically weighted regression to model spatial variations in the relationship between NDVI and climate factors (temperature and precipitation) across the Serengeti district. The GWR model suggests that the climate variables (temperature and precipitation) and elevation could explain 68% of the total variance in NDVI across the entire Serengeti district, with an average local *R*^2^ of 0.68. It should be noted that local *R*^2^ values vary substantially across the study area, indicating that the effectiveness of these factors in explaining NDVI variations differs in different regions. Climate variables could account for up to 80% of NDVI variation in areas with lower elevations (Figure B8c), particularly in the eastern region. However, the western region, dominated primarily by grazing and cropping activities, showed lower local *R*^2^values, suggesting that other factors, such as land use, could be more influential in these areas, thus contributing to the lower explanatory power of climate variables. Most of the temperature coefficients were negative, except in the southeast region, where they were positive. Conversely, most precipitation coefficients (92%) were positive, suggesting higher precipitation levels were associated with higher NDVI values.

In general, the entire Serengeti district is expected to become warmer and wetter over time, with the magnitude of these changes being more remarkable in the long term (late century) under the SSP585 high-emission scenario. We anticipate a steady increase in mean temperature throughout the twentieth century. This warming trend is likely to have a considerable effect on the health and distribution of vegetation.

**Fig. B13:**
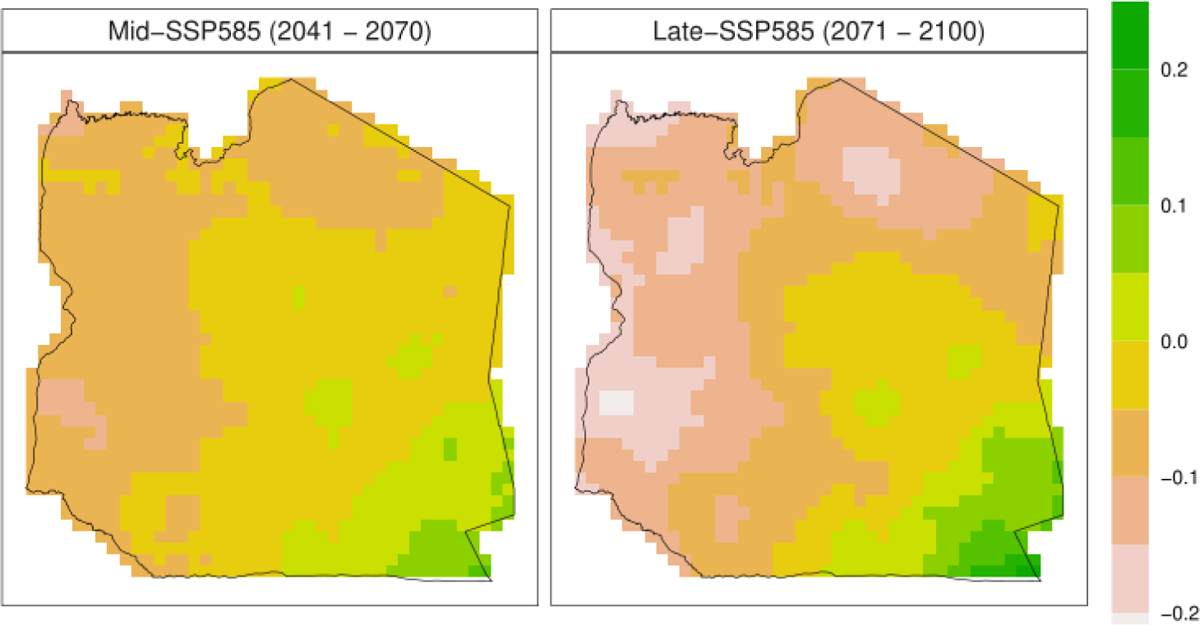
Relative changes in future NDVI predicted for mid (2041-2070) and late (2071-2100) centuries.

By combining climate projections and the GWR model, we predicted future NDVI values in the Serengeti. Our results indicate that, in general, NDVI will gradually decrease over time, with a 7.4 decrease by 2100. However, this trend is not uniform across the Serengeti district, with the western and northern parts of the region experiencing a decline in NDVI. In contrast, the southeastern region may see an increase. This could significantly impact livestock farmers who rely on these areas for grazing. This study has important implications for the Serengeti region. As temperatures rise and precipitation patterns change, vegetation dynamics could affect livestock grazing patterns. A decrease in NDVI values could lead to a reduction in the availability of grazing lands. This could result in competition for scarce resources, potentially altering contact patterns between livestock populations and increasing the risk of disease transmission. In addition, increased precipitation, particularly in the late century, may create favourable conditions for the spread of vector-borne diseases, posing risks to livestock and human health. These projected changes in temperature, precipitation, and NDVI could have a negative impact on the livelihoods of agropastoralist households who rely on livestock herding and agriculture. Localised approaches, effective adaptive tactics, and resource management will be necessary to address these challenges in the face of climate-driven ecological transformations.

We have described the complexity and spatial variability of the relationship between climate and vegetation in the Serengeti region. However, there are some potential drawbacks to this research. Our study assumes that the observed relationship between vegetation and climate will remain constant and that the contribution of other factors, as represented by model residuals, will remain unchanged over time. We did not consider other ecological variables, such as soil properties, which can impact ecosystem dynamics. Furthermore, our model does not consider the anthropogenic impacts on vegetation, including the complexities associated with predicting the effects of factors such as increasing population, economic development, and land management practices, which may have different results for vegetation growth and distribution.

## Data availability

The dataset used for this analysis records the geographical coordinates of villages and resource points, together with the monthly inter-village connections on shared resources in the Serengeti district. This dataset belongs to Divine Ekwem, and we do not have permission to share it publicly. Please contact the corresponding author for information on accessing this data.

## Code availability

All codes used in this manuscript were developed in R and are available in a private GitHub repository at https://git.ecdf.ed.ac.uk/s2120283/resource driven movement. Access will be granted upon request from the corresponding author.

## Authors’ contributions

TS, DE, PJ, JE, SC, and RK designed the study. TS developed the code, performed the analyses, produced visualisations, and wrote the original draft of the manuscript. DE, DF, and PP collected and shared the data. All authors read, reviewed, and approved the submitted version.

